# Fish oil Nano-emulsion Kills Macrophage: Ferroptosis and Autophagy Triggered by Catalase-catalysed Superoxide Eruption

**DOI:** 10.1101/2022.02.02.478786

**Authors:** Guanzhen Gao, Jingru Zhou, Huiqin Wang, Lijing Ke, Jianwu Zhou, Yanan Ding, Wei Ding, Suyun Zhang, Pingfan Rao

## Abstract

Fish oil is increasingly utilized in the form of nano-emulsion as nutrient and function fortifier. The nano-emulsion’s high polyunsaturated fatty acids content and electron donors at oil/water interface could be a potential site of redox reaction, if a previously unrecognised trigger was provided. Here we report that a vigorous superoxide production occurred in fish oil nano-emulsion in the presence of mammalian catalase in both acellular and cellular systems. Furthermore, the resulting superoxide increased cytosolic ROS and membrane lipid peroxidation of murine macrophage, and eventually caused fatal oxidative damage, which involves autophagy and ferroptosis but not apoptosis. The cell death was significantly inhibited by a catalase-specific inhibitor. The ferroptosis was independent of protease Caspase-3 activation or glutathione peroxidase suppression. Our findings discover a hidden risk factor of the widely acclaimed fish oil emulsion and suggest a novel mechanism of cellular damage by dietary lipids on mucosal layer of the alimentary tract.

## INTRODUCTION

Fish oils are highly rich in long-chain *ω*-3 polyunsaturated fatty acids (PUFAs), which contain a double carbon-carbon bond at the third carbon atom (*n*-3 position) from the methyl end of the carbon chain, consisting of docosahexaenoic acid (DHA; 22:6) and eicosapentaenoic acid (EPA; 20:5)^1^. With mounting evidence demonstrating various benefits of DHA and EPA for brain development, cardiovascular health^2^, and inflammation^3^, fish oil has found its wide applications in food and beverage products as nutrient fortifier^4^, and in parenteral nutrition formula to provide extra physiological benefits beyond its basic nutrition and energy^5^.

Fish oil has poor palatability and high susceptibility to oxidation. To overcome this problems, delivery systems based on oil-in-water emulsion or biopolymer complexes^6,7^ were employed for its applications in food and other health products^8^. Among these, fish oil nano-emulsion (FONE) emerges as a promising solution.

Fish oil emulsion has been used in bakery products, sauces, dressing, dairy products, beverages, and supplements^9^. It is proven to be beneficial for patients suffering from inflammatory bowel disease, rheumatoid arthritis, and asthmatics^10^. Various types of emulsifiers have been employed to enhance the stability of fish oils emulsions, including small molecule surfactants (polysorbate or sorbitan), amphiphilic polysaccharides (gum arabic or modified starch), phospholipids (soy, egg, or dairy lecithin), and amphiphilic proteins (caseinate or whey protein isolate)^11,12^. The poor oxidative stability of fish oil emulsion is commonly addressed by adding small molecular weight antioxidants such as butylated hydroxyanisole, butylated hydroxytoluene, *et al*. Furthermore, some antioxidant enzymes, i.e., superoxide dismutase (SOD, EC 1.15.1.1), have been applied to successfully prevent lipid oxidation^13^ in emulsions.

Catalase (CAT) is another crucial antioxidant enzyme present in almost all aerobic organisms. It catalyses hydrogen peroxide into water and molecular oxygen^14^, or oxidises low molecular weight alcohols in the presence of low concentrations of hydrogen peroxide (peroxidase activity)^15^. However, catalase did not prevent the lipid oxidation of polysorbate 80-stabilized fish oil nano-emulsion while induced rapid demulsification. The demulsification was found to be irrelevant of the lipid peroxidation, but closely correlated to catalase’s enzymatic activity^16^. Although its mechanism remains an enigma, the demulsification has been hypothesized to correlate to hydroperoxyl radicals on the oil/water interface rather than the catalase-H_2_O_2_ reaction^16^. The accelerated lipid peroxidation indicates catalase can induce the generation of reactive oxygen species (ROS) in the fish oil emulsion. It is in good agreement with the earlier report that mammalian catalase possesses oxidase activity in the presence of oxygen and electron donor substrates and in the absence of H_2_O_2_. It has been predicted that superoxide could potentially be formed as the intermediate product of such oxidation^17^.

It is interesting to identify an unexpected new factor, catalase, determining the physicochemical stability of fish oil emulsion. And yet it is far more urgent and critical to gain a full-picture of the catalase-triggered changes in cytosolic redox homeostasis, molecular packing of biological membrane lipids, and cell viability particularly when it occurs intracellularly. Like most nanoparticles, fish oil nano-droplets are expected to be engulfed by macrophages and internalized into peroxisomes^18,19^. As a key regulator in overall cellular lipid and ROS metabolism^20^, peroxisomes are highly abundant in catalase^21^. Subsequently, the catalase-triggered demulsification and the accompanying ROS generation would be readily to occur inside peroxisomes. When fish oil emulsions are increasingly utilized in food and medicine, it is imperative to clarify concerns with even the slimmest implication of risk to health.

In the present studies, we aim to elucidate the catalase-mediated redox reaction in the oil-in-water nano-emulsions of fish oil and polysorbate 80, which may accompany or result in the generation of superoxide, and to investigate the biological impacts of such reaction when occurring outside or inside the macrophages, including its influences on cell viability, cytosolic ROS and membrane lipid peroxidation level, and the possible causes of mediators of such changes.

## MATERIALS AND METHODS

### Materials

Dulbecco’s Modified Eagle Medium (DMEM), penicillin-streptomycin, Phosphate-buffered saline (PBS), 5-(and-6)-chloromethyl-2’,7’-dichlorodihydrofluorescein diacetate, acetyl ester (CM-H_2_DCFDA), C11-BODIPY™ 581/591, Hoechst 33342, Halt Protease and Phosphatase Inhibitor Cocktail (100×), and Tissue Protein Extraction Reagent (T-PER) were purchased from Thermo Fisher Scientific, Inc. (Waltham, MA, USA). Fetal bovine serum (FBS) was purchased from Bovogen Biologicals Pty Co., Ltd (Keilor, Victoria, Australia). Polysorbate 80, dimethyl sulfoxide (DMSO), catalase from bovine liver, superoxide dismutase from bovine erythrocytes, and 3-Amino-1,2,4-triazole (3-AT) (≥95% TLC) were purchased from Sigma-Aldrich (St. Louis, MO, USA). Z-VAD-FMK, Necrostatin-1, Liprostatin-1, and 3-(4,5-dimethylthia-zol-2-yl)-3,5-diphenylformazan (MTT) (≥99.59%) was purchased from Med Chem Express (Monmouth Junction, NJ, USA). Malondialdehyde (MDA) assay kits, lactate dehydrogenase (LDH) cytotoxicity assay kits, Catalase assay kits, and Annexin V-FITC/PI apoptosis assay kits were purchased from Beyotime Biotechnology Co., Ltd. (Shanghai, China). Two kinds of fish oil containing 30 % (mode: Omega-3 18/12TG, Lot: J010VN20180612P1812) and 70% (mode: Omega-3 70/10EE, Lot J002QD20190424P7010) of PUFAs, respectively was kindly provided by Sinomega Biotech Engineering Co., Ltd (Zhoushan, Zhejiang, China). 2-methyl-6-p-methoxyphenylethynyl-imidazopyrazinone (MPEC), a chemiluminescence probe specifically reacted with superoxide, was purchased from Atto Co., Ltd. (Tokyo, Japan). Anti-microtubule-associated protein 1 light chain 3B (LC3B) primary antibodies (Catalog NO: 4108), anti-cleaved Caspase 3 primary antibodies (Catalog NO: 9661), anti-mixed lineage kinase domain-like protein (MLKL) primary antibodies (Catalog NO: 37705), and anti-phospho-MLKL primary antibodies (Catalog NO: 37333) were purchased from CellSignaling Technology, Inc. (Danvers, MA, USA). Anti-Pro-Caspase 3 primary antibodies (Catalog NO: ab90437), anti-glutathione peroxidase 4 (GPX4) primary antibodies (Catalog NO: ab125066), and anti-glyceraldehyde 3-phosphate dehydrogenase (GAPDH) primary antibodies (Catalog NO: ab181602) were purchased from Abcam (Cambridge, MA, USA). Other chemicals were of analytical grade and used without further purification.

### Fish oil nano-emulsions preparation

Two kinds of fish oil nano-emulsions were prepared according to our previous report^16^, one was prepared with the fish oil containing more than 30% omega-3 PUFAs named FONE30, and another was prepared with fish oil containing more than 70% omega-3 PUFAs named FONE70. Briefly, Polysorbate 80 (50 μL) was dissolved in 50 mL deionized water (Milli-Q, Millipore, U.S.) with stirring at 40°C. Fish oil (250 μL, 5‰ v/v) was controlled to slowly drip into polysorbate solution with constant stirring, then homogenized at 18,000 rpm for 5 min (high-speed homogenizer T18DS25, IKA, Staufen, Germany) and filtrated with a 0.22 μm polyethersulfone (PES) membrane filter (Millipore, USA) in 40°C water bath to obtain the fish oil emulsion.

### Characterization of fish oil nano-emulsions

The particle size and ζ-potential of fish oil emulsions were measured at room temperature using a ZetaSizer Nano-ZS instrument (Malvern Instruments Co. Ltd., Worcestershire, UK). Each measurement was repeated three times. The emulsions might be diluted to achieve appropriate light scattering intensity. The fish oil emulsions were subjected to the 300-mesh copper grid, stained with 1% (w/v%) of uranyl acetate, and then observed with transmission electron microscopy (TEM, Joel JEM-1230, Tokyo, Japan) at an acceleration voltage of 80 kV.

### Measurement of ROS and hydroperoxide content in FONE initiated by catalase

MPEC probes can detect superoxide in aqueous or biological samples with a higher sensitivity than MCLA or Luminol^22^. MPEC probes were used here to detect the superoxide produced in the fish oil emulsion, in the absence or presence of catalase (0, 1,500, 3,000, 4,500, 6,000, 10,000 U/mL). ROS in 5 mg/mL FONE30 or FONE70 was determined using 10 μM MPEC (Atto, Tokyo, Japan). The SOD (2,000 U/mL) was added into the emulsion in the presence of catalase (10,000 U/mL) to verify the type of free radical. Luminescence was measured using an AB-2280 Luminescencer NIR system (Atto, Tokyo, Japan). Each sample was determined with at least three replicates.

Hydroperoxide content in FONE was determined based on ferrous oxidation–xylenol orange (FOX) method^23^. Briefly, samples were mixed with 100 μL methanol in the test tubes with a vortex mixer for 3-6 s, then 900 μL FOX reaction mixtures (25 mM H_2_SO4, 100 μM xylenol orange, 250 μM FeSO4, and 4 mM BHT in 90% (v/v) methanol) was added and incubated for 30 min at room temperature. The absorbance was determined at 560 nm.

### Cell viability assay

Murine macrophage (Raw 264.7) was kindly provided by Stem Cell Bank, Chinese Academy of Sciences (Shanghai, China). Raw 264.7 cells were cultured in DMEM supplemented with 10% FBS, 100 U/ml penicillin, and 100 μg/ml streptomycin at 37◻ with 5% CO_2_ 95% humidified incubator. The cells (2×10^4^ cells/mL) were seeded into 96-well plates. Cell viability was evaluated with MTT assay and LDH cytotoxicity assay. The effect of FONE alone on cell viability was measured at the lipid concentrations of 0, 16.7, and 20 μg/mL. The effect of FONE-exogenous catalase interaction on cell viability was determined at 16.7 μg/mL FONE70 in the presence of exogenous catalase at 5 and 15 U/mL, respectively. The influences of endogenous catalase of macrophages on the cells’ viability was investigated at 20 μg/mL FONE70. An endogenous catalase inhibitor, 3-Amino-1,2,4-triazole (3-AT) (0.2 and 2 mM), was used to verify the effects of endogenous catalase. The types of cell death were identified at 16.7 μg/mL FONE70 with/without 15 U/mL exogenous catalase, in the presence or absence of either Z-VAD-FMK (apoptosis inhibitor, 20 μM), or Necrostatin-1 (necroptosis inhibitor, 40 μM), or Liprostatin-1 (ferroptosis inhibitor, 1 μM). The influences of catalase, 3-AT, and polysorbate 80 on cell viability were evaluated, too, for comparison.

### Measurement of total ROS and lipid peroxidation levels

Total ROS and lipid peroxidation levels were determined using fluorogenic probes of 5-(and-6)-chloromethyl-2′,7′-dichlorodihydrofluorescein diacetate, acetyl ester (CM-H_2_DCFDA) and C11-BODIPY™ 581/591, respectively^24^. After incubation with the emulsion, CAT, or 3-AT for 2 h, murine macrophages were loaded with CM-H_2_DCFDA (10◻μM) or C11-BODIPY 581/591 (5 μM) for 20 min at 37°C in serum-free DMEM medium. After incubation, the medium containing CM-H_2_DCFH-DA was removed. CM-DCF fluorescence inside cells was measured at 485 nm for excitation and 538 nm for emission with FlexStation® 3 Multi-Mode Microplate Reader (Molecular Devices, Sunnyvale, CA, USA). The amount of emitted fluorescence was correlated to the quantity of reactive oxygen species (ROS). Fluorescence of C11-BODIPY™ 581/591 was measured by simultaneous acquisition of the green (484/510 nm, oxidized) and red signals (581/610 nm, reduced) with FlexStation® 3 Multi-Mode Microplate Reader (Molecular Devices, Sunnyvale, CA), and the ratio of green/red fluorescence was determined as a measure of oxidation of C11-BODIPY™ 581/591.

Samples for fluorescent imaging of lipid peroxidation observation were performed with a Leica TCS SP8 Laser Scanning Confocal Microscope (Leica microsystems, Mannheim, Germany). The fluorescent signals of the non-oxidized and oxidized forms were acquired using simultaneous excitation. The non-oxidized form of the dye was excited at 561 nm, and signals were detected at 570–660 nm. The oxidized form of the probe was excited at 488 nm, signals were detected at 500 to 560 nm. The whole process was repeated twice to confirm the reproducibility.

### Assay of intracellular contents of MDA and catalase

The concentration of MDA and the activity of catalase were all determined by using commercially available kits according to the manufacturer’s instructions. The catalase activity of cells treated with 3-AT was detected with the Catalase Assay Kit. The activities of enzymes were expressed as units per milligram protein. The MDA of cells treated with fish oil nano-emulsion present/absent of 3-AT or catalase were measured at a wavelength of 532 nm by reacting with thiobarbituric acid (TBA) to form a stable chromophoric production. Values of MDA level were expressed as nanomoles per milligram protein.

### Flow cytometry assay

Annexin V-FITC/propidium iodide (PI) double staining assay was used to measure apoptosis and necrosis of the RAW 264.7 cells. Cultured RAW 264.7 treated with 16.7 μg/mL FONE with catalase (0, 5 and 15 U/mL), or FONE70 (0, 16.7 and 20 μg/mL), or 20 μg/mL FONE70 with 3-AT (0, 5, and 15 U/mL) for 24 h were collected. After washing the cells with phosphate-buffered saline (PBS) three times, each group of cells was stained following the manufacturer’s instructions. The number of apoptotic and necrosis cells was detected by flow cytometry (EasyCell206A1, Wellgrow Technology Co., Ltd., Shenzhen, China) and analysed using FlowJo software. Each group was repeatedly measured three times.

### Autophagy Flux Detection

The Cyto-ID fluorescent reagent specifically labels autophagic vacuoles and colocalizes with LC3^25^. Thus, autophagosome formation in cells was detected using the Cyto⋻ID Autophagy Detection Kit (ENZO Life Sciences, Inc., Farmingdale, NY, USA). The RAW 264.7 cells were treated with FONE (0, 16.7, and 20 μg/mL), with catalase (0, 5, and 15 U/mL) or 3-AT (0, 5, and 15 U/mL) for 24 h and then subjected for the Cyto-ID autophagy detection. The assay was performed according to the manufacturer’s instructions with FlexStation® 3 Multi-Mode Microplate Reader (Molecular Devices, Sunnyvale, CA, USA). The Cyto-ID fluorescence was determined at Ex 480 nm/Em 530 nm, while Hoechst 33342 fluorescence was determined at Ex 340 nm/Em 480 nm. The latter was used to normalize Cyto-ID fluorescence to indicate the autophagy flux.

### Western blotting analysis

RAW 264.7 cells were treated with fish oil emulsion in the presence or absence of either 2.0 mM 3-AT or 15 U/mL catalase for 24 h. The cells were washed twice with ice-cold PBS and lysed with T-PER tissue protein extraction reagent containing the ‘Halt’ protease and phosphatase inhibitor cocktail. The lysates were centrifuged at 14,000 × g for 15 min at 4°C. The protein concentration of each sample was measured using a bicinchoninic acid protein assay. Equal amounts (60 μg) of protein were separated using 10% sodium dodecyl sulfate-polyacrylamide gel electrophoresis (SDS-PAGE) and then transferred to polyvinylidene fluoride membranes. The membranes were blocked with Tris-buffer saline containing polysorbate 20 (TBST) containing 5% BSA for 1 h at room temperature, then washed three times (5 min each) with Tris-buffer saline containing polysorbate 20 (TBST) and incubated with the respective primary antibodies: anti-GAPDH antibody (Abcam, 1:10000), anti-GPX4 antibody (Abcam, 1:10000), anti-Pro-Caspase 3 (Abcam, 1:1000), anti-Cleaved Caspase 3 antibody (CellSignaling Technology, 1:1000), anti-LC3B antibody (CellSignaling Technology, 1:1000), and anti-MLKL/phospho-MLKL antibody (CellSignaling Technology, 1:1000) overnight at 4°C. After primary antibody incubation, followed by HRP-conjugated secondary antibodies, a signal was developed by SuperSignal® West Dura Extended Duration Substrate (Thermo Fisher Scientific, Waltham, MA, USA). Western blotting experiments were repeated three times, and the quantitative analysis of western blotting results was performed with Image J1.47 (National Institutes of Health, Bethesda, MD, USA).

### Statistical analysis

Statistical analysis was performed using the GraphPad Prism software (La Jolla, CA, USA). Data are expressed as mean ± standard error of the mean (SEM) (*n* = 3-6). Using one-way ANOVA followed by the Tukey’s *t*-test for post-hoc analysis to compare the data of different experimental groups, *P* < 0.05 was considered to show statistically significant differences.

## RESULTS

### Fish oil nano-emulsions

The two kinds of fish oil, e.g., F30 and F70, used here have a high content of *ω*-3 and *ω*-6 PUFAs, which account for 50.0% and 93.9% of total lipids, respectively (as shown in Table S1). Their major difference was in the content of EPA.

Two fish oil nano-emulsions (FONE) were prepared with good reproducibility. The FONE droplets were characterized for their particle size and ζ-potential with dynamic light scattering methods. Their morphological profiles were observed with TEM. As is shown in Fig. 1A, B, C, and D, the particle size of FONE30 (*ω*-3 PUFAs > 30%) and FONE70 (*ω*-3 PUFAs > 70%) was 119.2 ± 0.3 nm and 128.7 ± 0.2 nm, while the ζ-potential was −16.4 ± 0.1 mV and −16.7 ± 0.3 mV, respectively. FONE70’s nanoparticle and ζ-potential were both slightly higher than FONE30’s. Both emulsions displayed nano-droplets with similar size distribution in TEM observation, as shown in Fig. 1E and 1F.

**Fig. 1.**
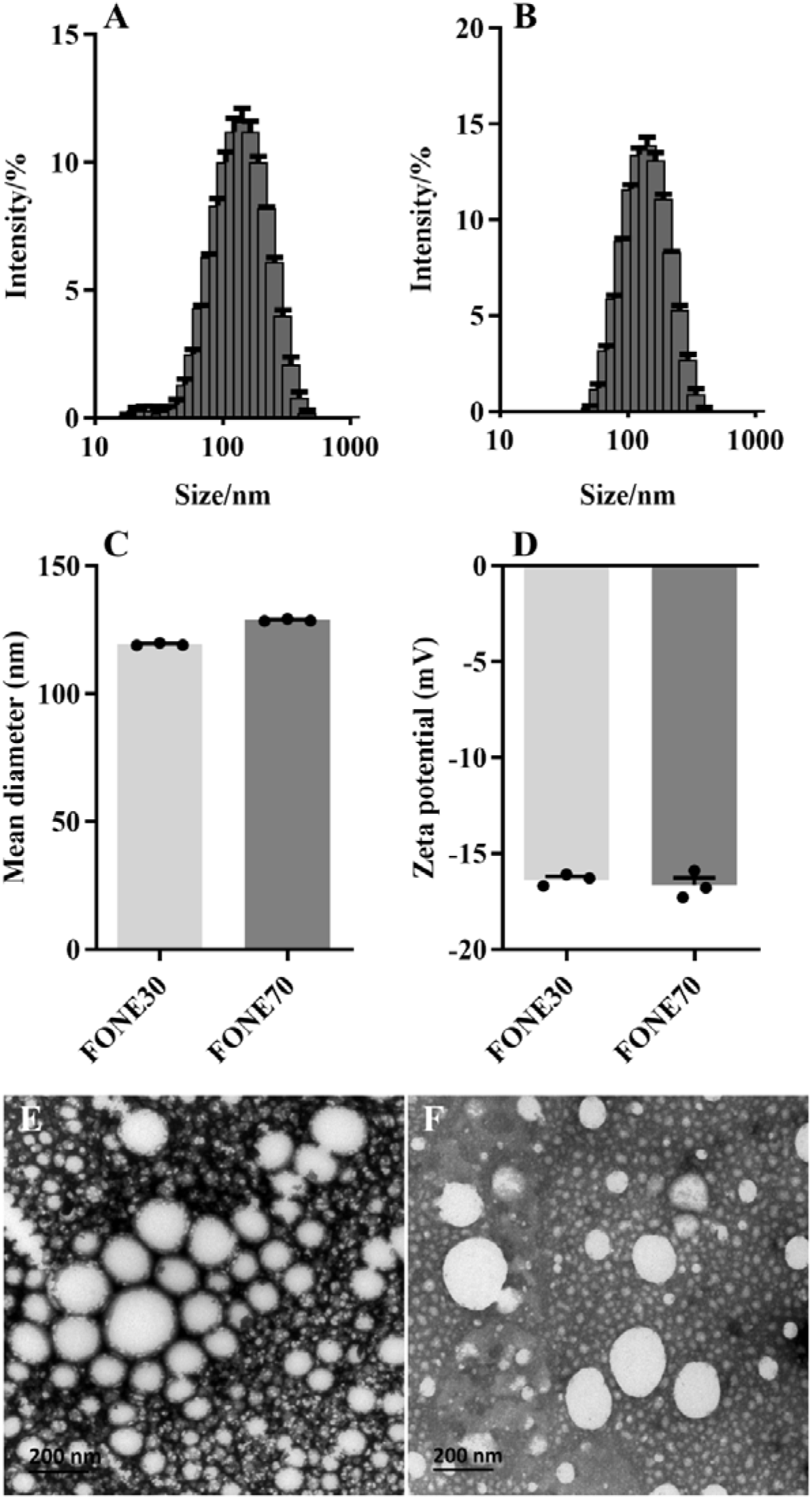
Granular properties of oil-in-water fish oil nano-emulsions (FONE) (A) size distribution of FONE30; (B) size distribution of FONE70; (C) mean diameter of FONE30 and FONE70; (D) Zeta potential of FONE30 and FONE70; Data were expressed as the mean ± SEM (*n* = 3). (E) TEM observation of FONE30; (F) TEM observation of FONE70.

### Catalase-triggered ROS eruption in FONE

The reactive oxygen species (ROS) generated in the fish oil nano-emulsions, either in the presence or absence of catalase, were detected and measured by using a luminescence-based assay. ROS determinations using luminescent probe MPEC were shown in Fig. 2A. Catalase boosted ROS by 10 folds in both FONE30 and FONE70. The ROS level increased in parallel with the rising catalase concentration and topped at 6000 U/mL of catalase. At this point, the amount of ROS is greater than the superoxide generated by 800 μM xanthine with 0.1 U xanthine oxidase (Supplemental Fig. S1). Notably, it is already far beyond the fatal dosage to mammal cells. It is known that ROS generated by 30 μM xanthine and 0.1 U xanthine oxidase was highly cytotoxic ^26^. Moreover, the catalase-induced ROS in FONE30 and FONE70 was quantitative in proportion to the PUFAs content of emulsions. As it produced a higher amount of ROS, FONE70 was used in the cellular biology studies of FONE and catalase on macrophages.

**Fig. 2.**
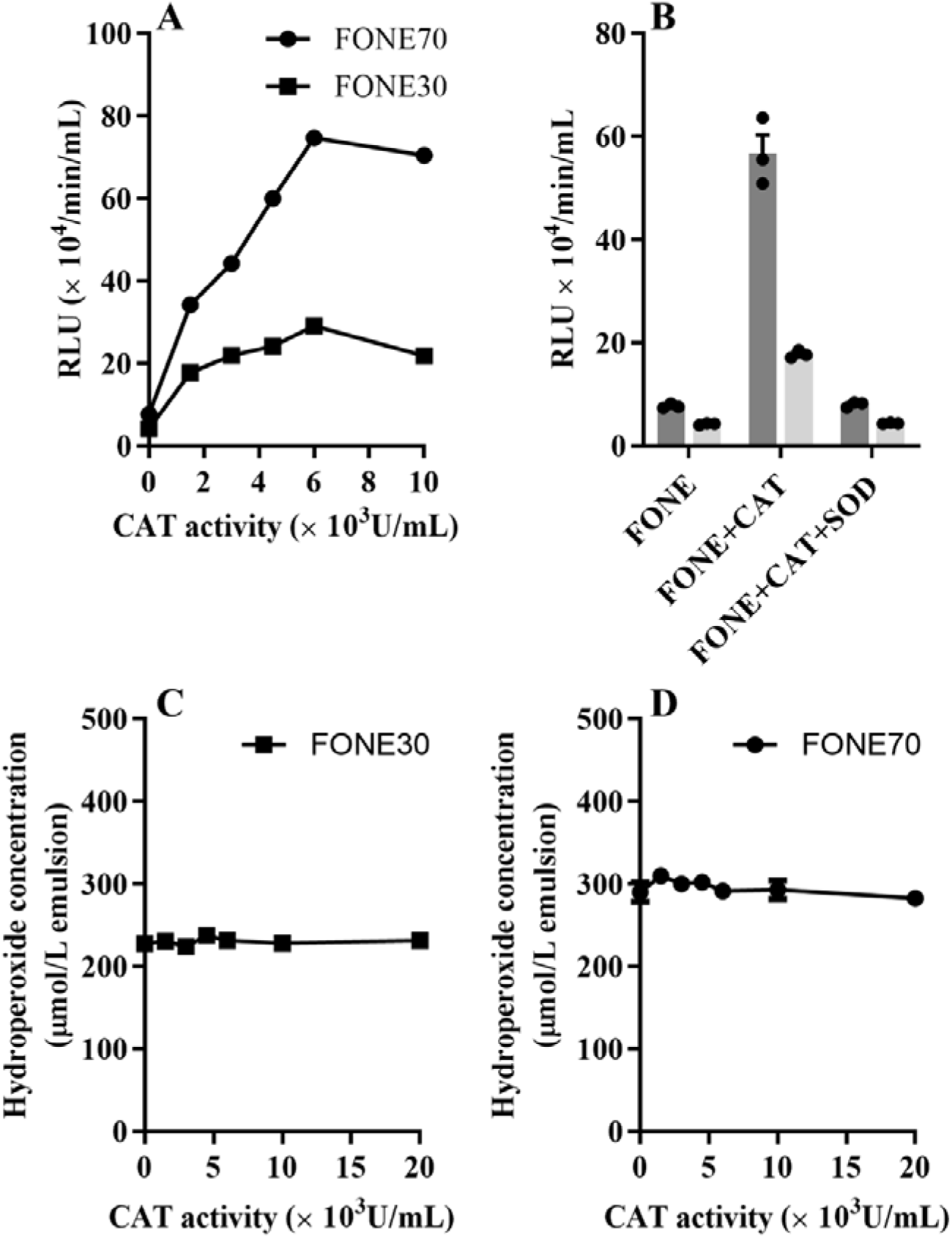
Superoxide generation and hydroperoxide content of FONE with/without CAT. (A) Superoxide induced MPEC luminescence in FONE30 or FONE70 in the presence of catalase; (B) superoxide induced MPEC luminescence in FONE30 or FONE70, in the presence of catalase (10,000 U/mL) or SOD (2,000 U/mL); (C) hydroperoxide content of FONE30 in the presence of catalase, using ferric-xylenol orange assay; (D) hydroperoxide content of FONE70 in the presence of catalase using ferric-xylenol orange assay. FONE30 or FONE70 at 5 mg/mL and catalase at 0 U/mL, 1,500 U/mL, 3,000 U/mL, 4,500 U/mL, 6,000 U/mL, and 10,000 U/mL were used in the tests. Data were expressed as the mean ± SEM (*n* = 3, 4).

The erupted ROS was identified as superoxide by two methods. Firstly, the luminescent probe employed is highly superoxide-specific, owing to chemiluminescence only occurring in the reaction with the superoxide ion^27^. Secondly, the ROS triggered by catalase was nearly wiped out by superoxide dismutase (as shown in Fig. 2B), which scavenges superoxide specifically. The presence of a prominent amount of superoxide often heralds oxidative stresses and severe malfunction upon neighbouring cells.

Hydroperoxides are produced in the initiation and progress of polyunsaturated fatty acids autooxidation, which could be triggered by superoxide radicals in the surrounding microenvironment and catalysed by the exerted ferrous iron. The hydroperoxide content in FONE was determined by the Fenton reaction. However, as shown in Fig. 2C and 2D, hydroperoxide was found to be rather low in FONE30 (~ 200 μM) and FONE70 (~ 300 μM), regardless of the presence of iron-containing CAT. The CAT-induced superoxide surge did not elevate hydroperoxide content in the emulsions, elucidates that the CAT-induced superoxide did not oxidise the PUFAs in FONEs, at least not immediately, and the heme iron in CAT did not catalyse the generation of hydroperoxide in the system.

### Cell death caused by FONE and catalase in murine macrophages

The superoxide eruption catalysed by CAT in the FONE can cause oxidative stresses and damages to the surrounding biological subjects. Unlike in the test tubes, the living cells often have significant storage of oxidoreductases, including catalase, and therefore allow such oxidative stress to occur readily, even in the absence of exogenous CAT. For example, the peroxisomes of macrophages, containing a large quantity of endogenous catalase^21^, may encounter the emulsion droplets once the latter makes its entry into the cells. Would lymphocytes (macrophages) suffer from the direct exposure to the superoxide surge generated by the interaction between lipid nano-droplets and catalase? To answer this question, FONE70 was added to RAW 264.7 macrophages in the presence of endogenous or exogenous catalase.

As shown in Fig. 3A, the cell viability was significantly compromised corresponding to FONE70 concentration, dropping to less than 10% at 25 μg/mL of FONE70. The TC_50_ (Toxic Concentration 50%) of FONE70 on the macrophages was 16.5 μg/mL.

**Fig. 3.**
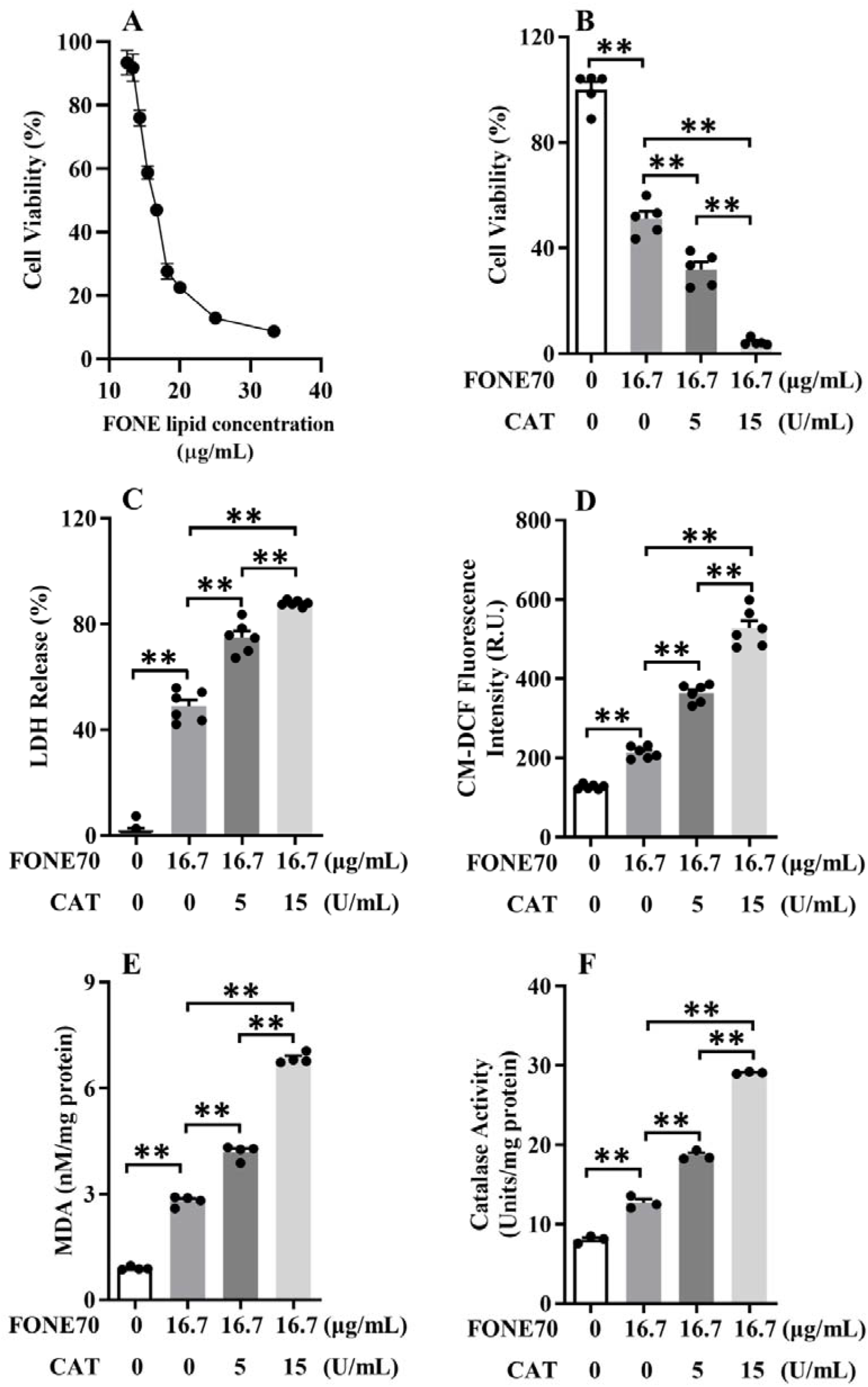
Effects of FONE70 and exogenous catalase on cell viability, intracellular ROS, and lipid peroxidation of Raw 264.7 cells. (A) Effects of FONE70 on cell viability, RAW 264.7 macrophages were treated with FONE70 at the lipid concentration of 12.5-33.3 μg/mL. The influences of catalase (0 U/mL, 5 U/mL, 15 U/mL) on (B) cell viability, (C) LDH release, (D) intracellular ROS quantified with CM-H_2_DCFDA fluorescent probe, (E) MDA content, and (F) catalase activity in cells treated with FONE70 at lipid concentration of 16.7 μg/mL. Data were expressed as the mean ± SEM (*n* = 3-6). Data were analysed by one-way analysis of variance (ANOVA) followed by the Tukey’s *t*-test for post-hoc analysis, \*\**P* < 0.01.

The drop in cell viability was dramatic at the oil concentrations lower than TC_50_ and was gradually attenuated at the concentrations higher than TC_50_.

The effects of FONE70 on intracellular ROS, LDH release, and MDA levels of macrophage were tested at the FONE70 concentration just above TC_50_ (16.7 μg/mL) and well above TC_50_ (20 μg/mL). The higher dose of fish oil emulsion caused a higher level of oxidation and more severe cell damage, as shown in Fig. S2. Briefly, FONE70 significantly increased the intracellular ROS (quantified by CM-DCF fluorescence) and MDA levels, promoted LDH release to extracellular space, and suppressed cell viability (MTT assay). The addition of exogenous catalase at 5 U/mL and 15 U/mL potentiated intracellular ROS, MDA, extracellular LDH, and lipid peroxidation simultaneously in a dose-dependent manner, although the exogenous CAT could not enter the cells and stimulated superoxide production extracellularly only. The FONE70 with 15 U/mL of exogenous catalase is highly toxic and destructive to the macrophages (as shown in Fig. 3B-E). Notably, the two CAT concentrations used in the cell tests were equivalent to the 1,500 U/mL CAT and 4,500 U/mL CAT used in the high lipid concentration FONE30/70 emulsions, in terms of consistent lipid/enzyme ratio. This choice of CAT concentration guarantees that the intensity of superoxide eruption is comparable between the cellular and acellular systems.

The fact that FONE70 alone has induced an excessive amount of ROS in cells and possessed potent cellular destructive power indicates the involvement of endogenous catalase. As shown in Fig. 3F, the endogenous catalase activity of the macrophage was increased from the intrinsic 8 U/mg protein to 13 U/mg protein after the 24-h incubation with FONE. The total CAT activity was lifted to 28 U/mg protein in the presence of exogenous CAT, which can be well explained by the amount of exogenous enzyme. It demonstrated that the fish oil emulsion could promote the expression of intracellular catalase, possibly as a response to the superoxide eruption when FONE droplets encountered the intrinsic CAT in cells.

To verify the involvement of endogenous CAT in FONE’s cellular impacts, a catalase-specific inhibitor, 3-amino-1,2,4-triazole (3-AT), was therefore employed to demonstrate the endogenous catalase’s causative role. 3-AT inhibits catalase activity by specifically and covalently modifying the enzyme and blocking the main access channel for oxidase substrates of catalase^28,29^. As shown in Fig. 4A, 3-AT significantly restored the viability of macrophage up to 80% in a dose-dependent manner, against 20 μg/mL FONE70. Consistently, 3-AT significantly counteracted the CAT’s stimulations on intracellular ROS, LDH release, and MDA, as is shown in Fig. 4B~D. The inhibition on endogenous catalase was verified by determining intracellular catalase activity, as shown in Fig. 4E. In addition, none of polysorbate 80, CAT, and 3-AT affected the cell viability, intracellular ROS, LDH release, and MDA levels of macrophage, when they were used separately (as shown in Fig. S3 & S4). It demonstrates that endogenous catalase can trigger the fish oil emulsion-induced ROS surge in macrophages, leading to cell damage and death.

**Fig. 4.**
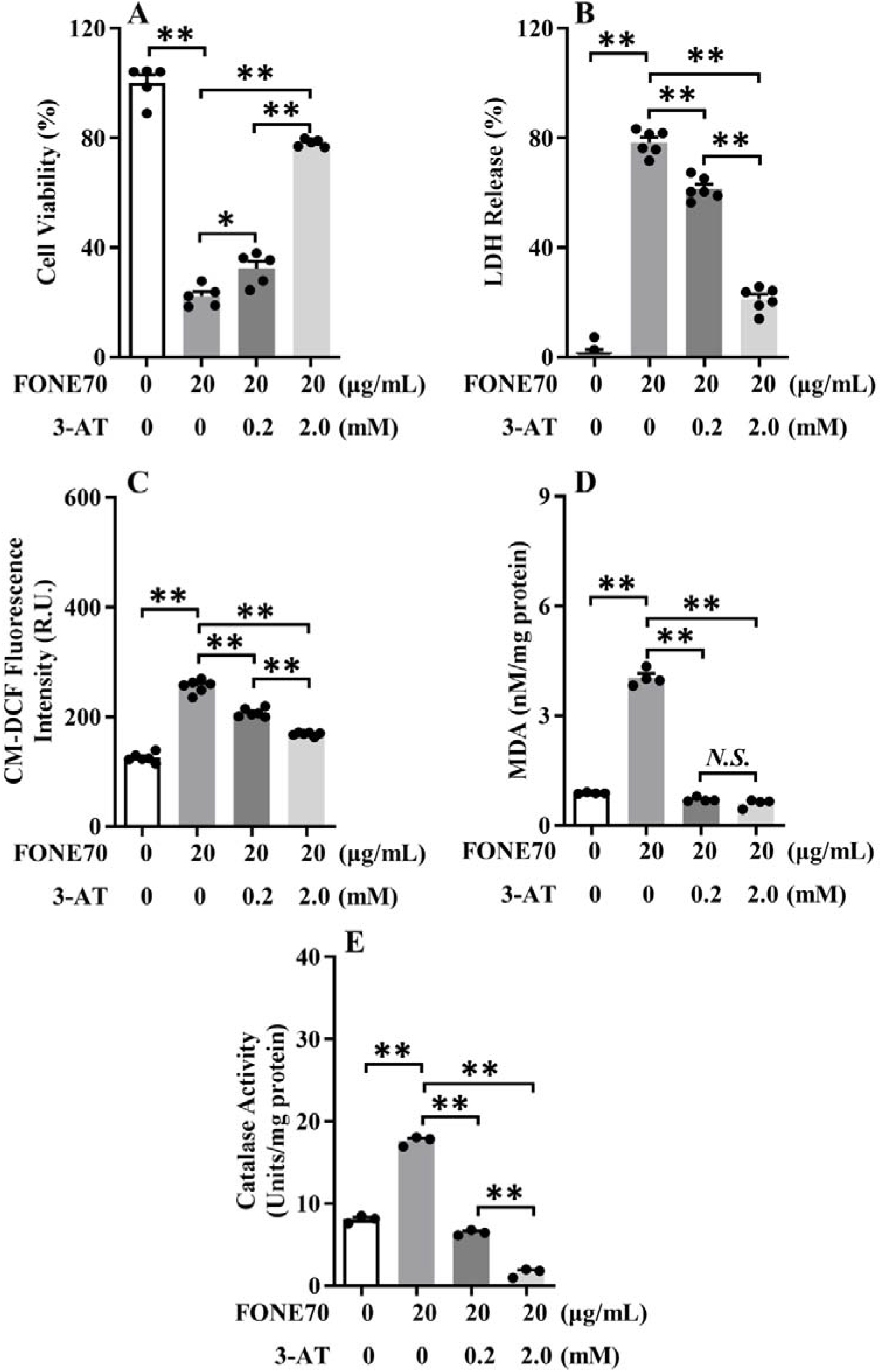
Inhibitory effects of catalase-inhibitor 3-AT on FONE-induced changes in Raw 264.7 cells. (A) Cell viability, (B) LDH release, (C) total ROS using a CM-H_2_DCFDA probe, (D) MDA content, (E) catalase activity in cells treated with FONE70 at lipid concentration of 20.0 μg/mL in the presence of catalase-inhibitor 3-AT (0 mM, 0.2 mM, 2.0 mM). Data were expressed as the mean ± SEM (*n* = 3-6). Data were analysed by one-way analysis of variance (ANOVA) followed by the Tukey’s *t*-test for post-hoc analysis, \**P* < 0.05, \*\**P* < 0.01.

### Identify the type of FONE-induced Cell Death

Intracellular oxidative stresses can be the reason behind different types of cell death, including necrosis and apoptosis. Necrosis refers to cell injury and unprogrammed death caused by external factors, such as toxin and infection, and is often associated with inflammatory responses. In contrast, apoptosis is programmed cell death, which is internally regulated by intrinsic signals and provides a protective mechanism against external factors and diseases^30^. As shown in Fig. 5A, 5B, and 5C, FONE70 induced necrosis which was further enhanced in the presence of catalase in a dose-dependent manner, while apoptosis remained unchanged except for an insignificant increase in the presence of 15 U/mL catalase (*P*>0.05). FONE70 at the higher dose alone (20 μg/mL) caused necrosis equivalent to the lower dose FONE70 (16.7 μg/mL) in combination with 5~15 U/mL exogenous catalase. Along with the abrogation of catalase by 3-AT, the FONE70 induced cell necrosis was fully prevented (Fig. 5D-5F). Neither CAT nor 3-AT induced necrosis or apoptosis when used alone (Supplemental Fig. S5). Therefore, the FONE-CAT triggered ROS eruption induced necrosis but not apoptosis.

**Fig. 5.**
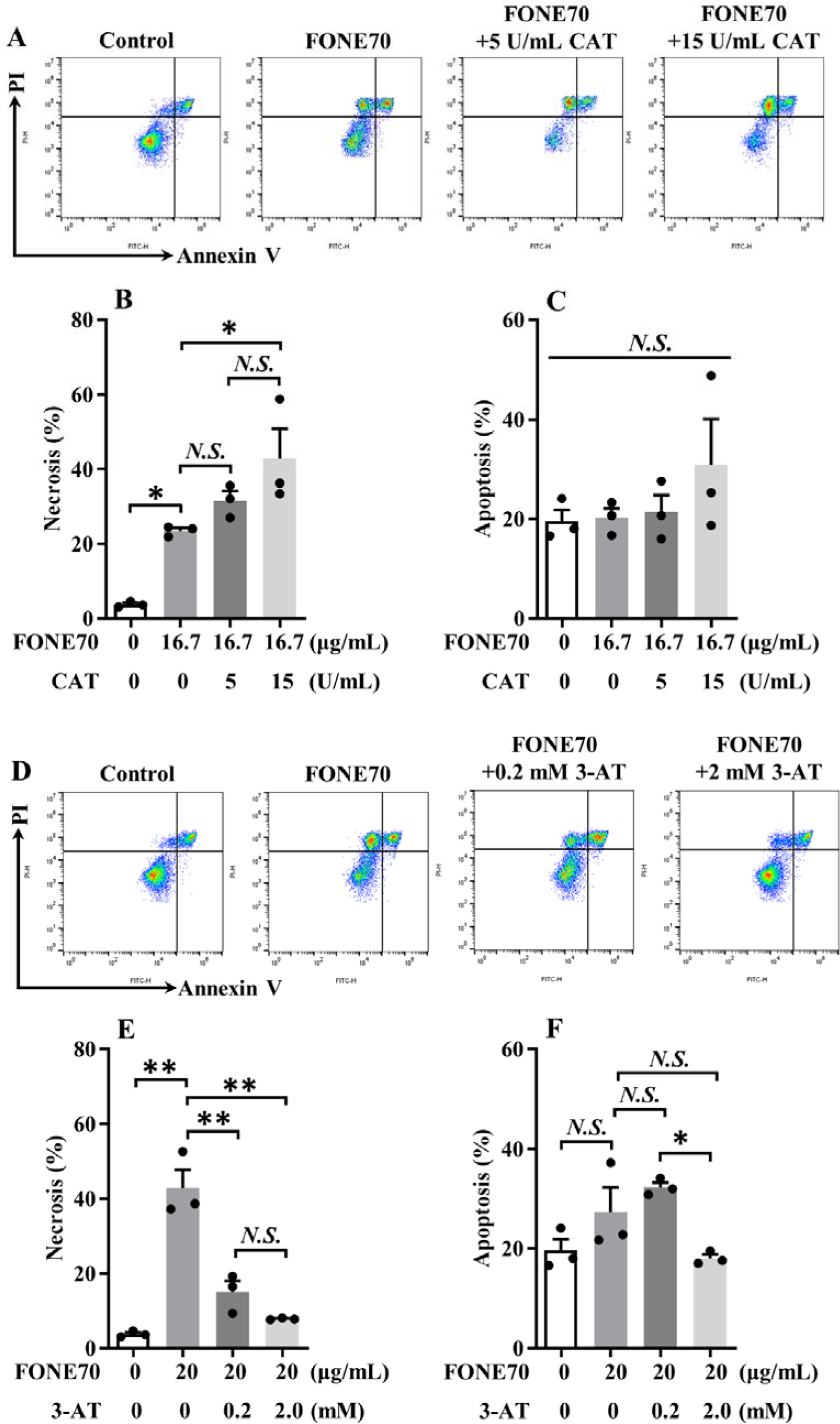
Flow cytometry analysis of FONE/catalase-induced cell death in macrophage Raw 264.7 cells. Cells were stained with Annexin V/propidium iodide (PI) double staining and detected by flow cytometry. Cells were treated with FONE70 at the concentration of 16.7 μg/mL in the presence of catalase (0 U/mL, 5 U/mL, 15 U/mL) for 24 h: (A) representative scatter plots of PI (y-axis) versus annexin V (x-axis); (B) the ratio of necrosis; (C) the ratio of apoptosis. Cells were treated with FONE70 at the concentration of 20.0 μg/mL in the presence of catalase-inhibitor 3-AT (0 mM, 0.2 mM, 2.0 mM) for 24 h: (D) representative scatter plots of PI (y-axis) versus annexin V (x-axis); (E) the ratio of necrosis; (F) the ratio of apoptosis. Data were expressed as the mean ± SEM (*n* = 3). Data were analysed by one-way analysis of variance (ANOVA) followed by the Tukey’s *t*-test for post-hoc analysis, \**P* < 0.05, \*\**P* < 0.01, *N.S*., no significance.

In order to gain more insights on the subtype of necrosis, three inhibitors were employed: Z-VAD-FMK as apoptosis inhibitor^31^, Necrostatin-1 as necroptosis inhibitor^32^, and Liprostatin-1 as ferroptosis inhibitor^33^. As shown in Fig. 6A, only the Liprostatin-1 effectively protected cells from fatal damage of FONE-induced ROS (*P*<0.01), while neither the caspase inhibitor Z-VAD-FMK nor receptor-interacting protein kinase 1 (RIPK1) phosphorylation inhibitor Necrostatin-1 affects the cell viabilities. It indicates that ferroptosis is possibly the underlying mechanism of cell death. The fact that CAT contains ferrous iron in the enzyme’s active centre adds weight to this explanation.

**Fig. 6.**
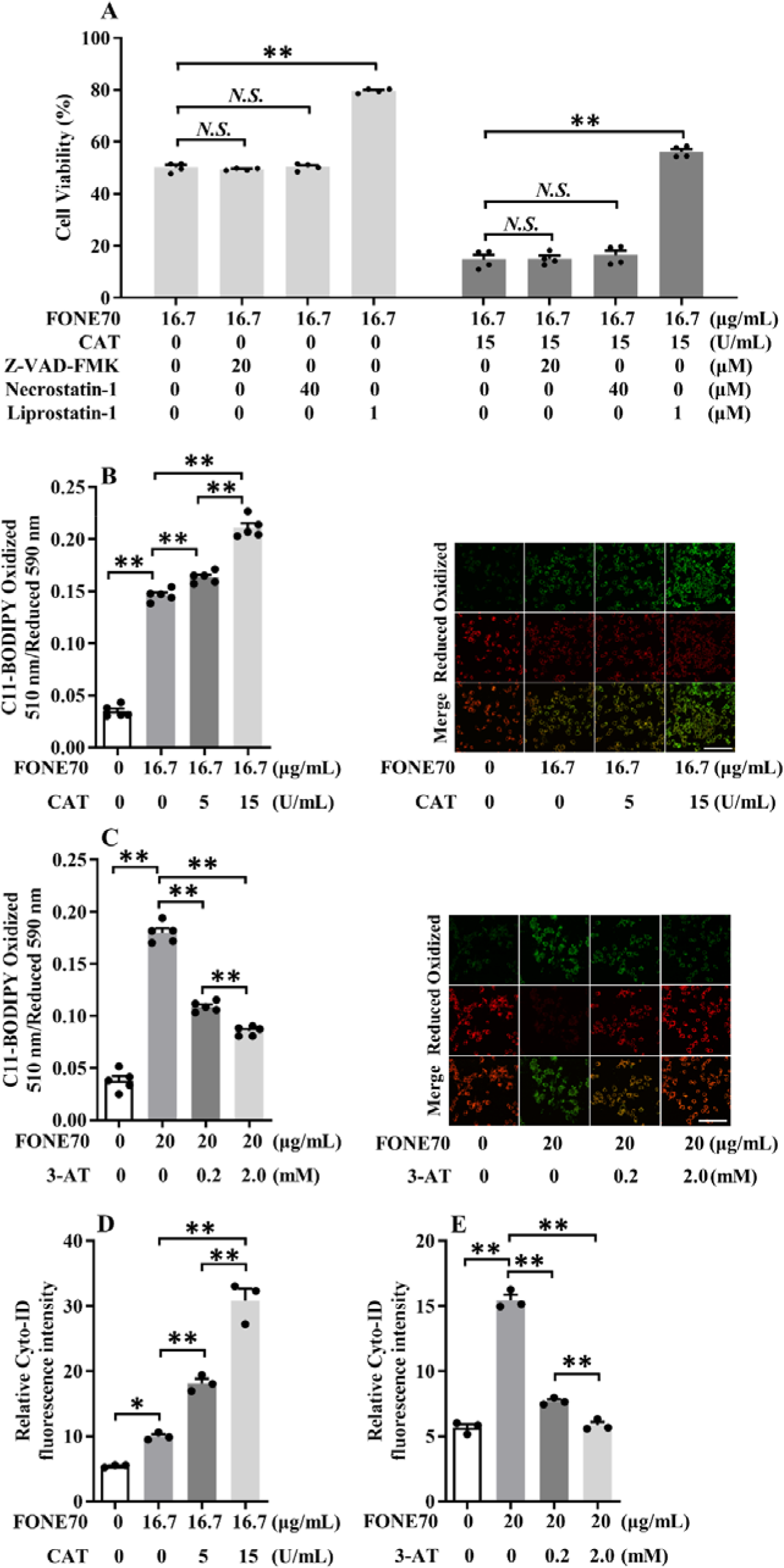
FONE70/catalase induced cell membrane lipid peroxidation, necrosis, and autophagy flux. (A) The effects of Z-VAD-FMK (apoptosis inhibitor), Necrostatin-1 (necroptosis inhibitor), and Liprostatin-1 (ferroptosis inhibitor) on the cell viability of FONE70 and FONE70-exogenous catalase interaction. (B) The ratio of oxidized 510 nm/reduced 590 nm and confocal microscopy images (scale bar = 100 μm) of lipid peroxidation using a C11-BODIPY probe and (D) autophagy flux in cells treated with FONE70 at the concentration of 16.7 μg/mL in the presence of catalase (0 U/mL, 5 U/mL, 15 U/mL). (C) The ratio of oxidized 510 nm/reduced 590 nm and confocal microscopy images (scale bar = 100 μm) of lipid peroxidation using a C11-BODIPY probe and (E) autophagy flux in cells treated with FONE70 at 20.0 μg/mL in the presence of catalase-inhibitor 3-AT (0 mM, 0.2 mM, 2.0 mM). Data were expressed as the mean ± SEM (*n* = 3 to 5). Data were analyzed by one-way analysis of variance (ANOVA) followed by the Tukey’s *t*-test for post-hoc analysis, \**P* < 0.05, \*\**P* < 0.01, *N.S*., no significance.

The cellular membrane lipid peroxidation is the typical consequence of oxidative stress and the signature event in ferroptosis. It could cause damage in cell membrane integrity and release of LDH, as observed earlier in this study. The peroxidation of cell membrane lipids of macrophages was accessed using a lipophilic fluorescent probe, C11-BODIPY, an indicator often used in ferroptosis studies^34^. Once the probe is oxidized by excessive ROS, the emitting green lights can be used to quantify lipid peroxides. It is quite clear in Fig. 6B that the FONE70 alone has already tripled the lipid peroxides content, while catalase significantly further intensified such peroxidation (*P* < 0.01). It demonstrates the aggressive membrane lipid peroxidation caused by the FONE, in the absence or presence of CAT, which may well be the direct cause of the cellular membrane destruction and eventual cell death. The catalase inhibitor 3-AT effectively suppressed the FONE-induced membrane lipid peroxidation (as shown in Fig. 6C), consistent with its protective effects on cell viability. Therefore, the FONE-induced macrophage necrosis is ferroptotic and catalase dependent.

On the other hand, autophagy is a catabolic process that involves recycling cytoplasmatic components and damaged organelles in response to diverse stress conditions, including starvation, pathogen infection, and toxic stress^35^. In recent years, oxidative stress has been proposed as the converging point of the above exogenous stimuli, as ROS serves as one of the primary intracellular signals sustaining autophagy^36^. Herein, the linkage between catalase-triggered ROS eruption and autophagy in the macrophages was examined with a rapid and quantitative assay hiring Cyto-ID fluorescence dye^25^. As shown in Fig. 6D&6E, the number of autophagosomes, indicated by Cyto-ID fluorescence, increased along with the addition of FONE, and was further fortified by exogenous catalase and inhibited by catalase inhibitor 3-AT. The sole use of CAT or 3-AT slightly increased autophagy flux, but in a much lower amount (Supplemental Fig. S5).

In summary, the FONE can induce intracellular superoxide eruption, lipid peroxidation, membrane destruction, and cause ferroptosis and autophagy of murine RAW 264.7 macrophage. The addition of exogenous catalase intensified such effects of FONE and catalase-specific inhibitor 3-AT inhibited them.

### Molecular pathway of FONE-induced ferroptosis and autophagy

To clarify the possible molecular pathway of ferroptosis and autophagy, the expression of cell death-related factors, e.g., LC3B-I, LC3B-II, Cleaved Caspase 3, Pro-Caspase 3, p-MLKL, MLKL, and GPX4, was determined in RAW 264.7 macrophages. The cells were treated by FONE70 in the absence/presence of exogenous catalase or catalase inhibitor 3-AT.

Autophagy protein LC3B is cleaved by autophagy-related 4 cysteine peptidase (ATG4) protease to form cytosolic LC3B-I. During autophagy, the cytosolic LC3B-I is converted to the autophagosome-associated LC3B-II, initiating the formation and lengthening of the autophagosome. Therefore, the ratio of LC3B-II/LC3B-I was commonly used as a marker for autophagy^37^. As shown in Fig. 7A-B, the ratio of LC3B-II/LC3B-I was significantly (*P* < 0.01) increased after treatment of FONE70 in a concentration-dependent manner. Exogenous catalase significantly enhanced such increase, while 3-AT completely inhibited it. It was in line with the results of autophagy flux, as shown in Fig. 6D&6E, confirming that cell death induced by FONE involved autophagy.

**Fig. 7.**
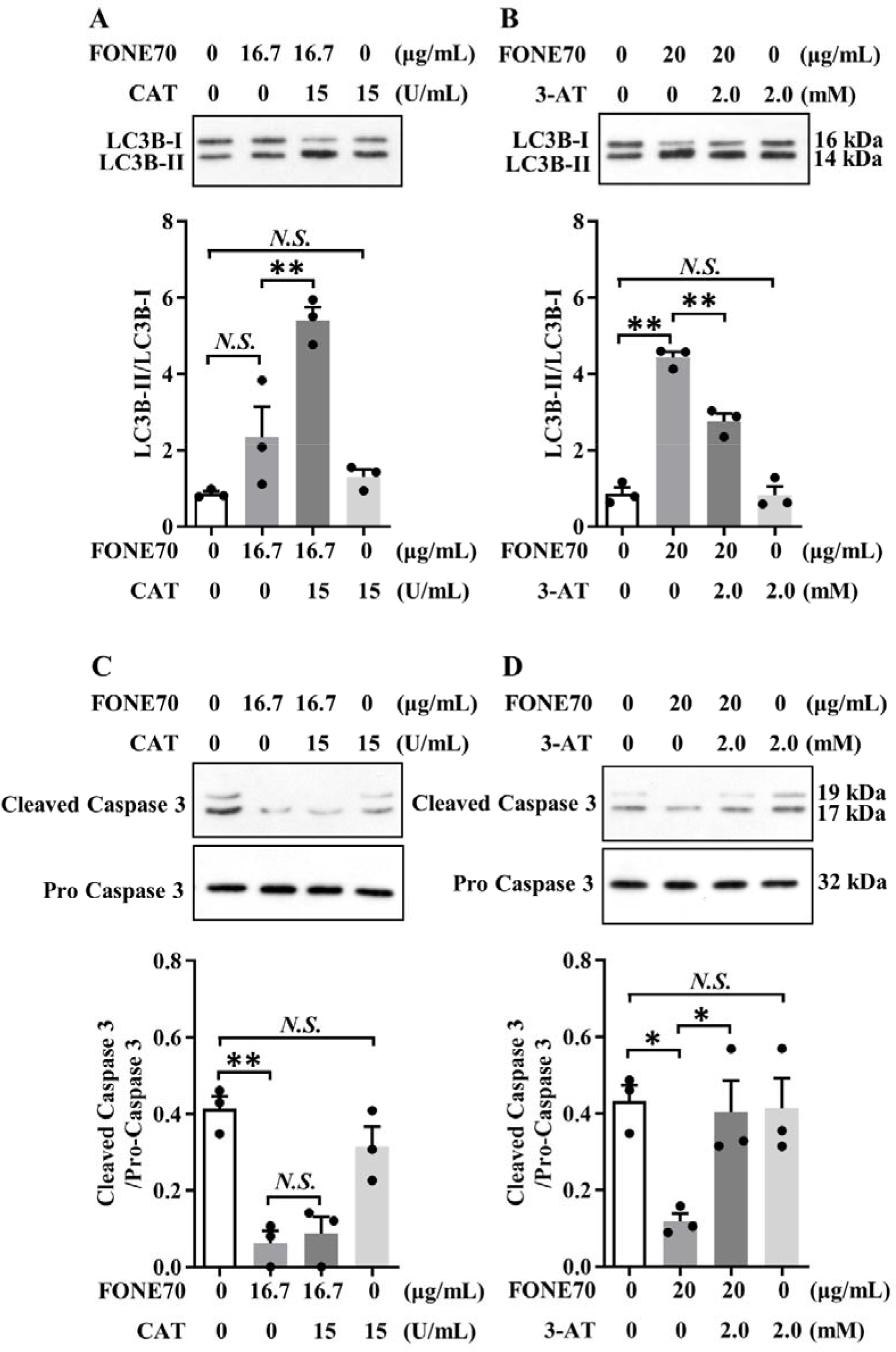
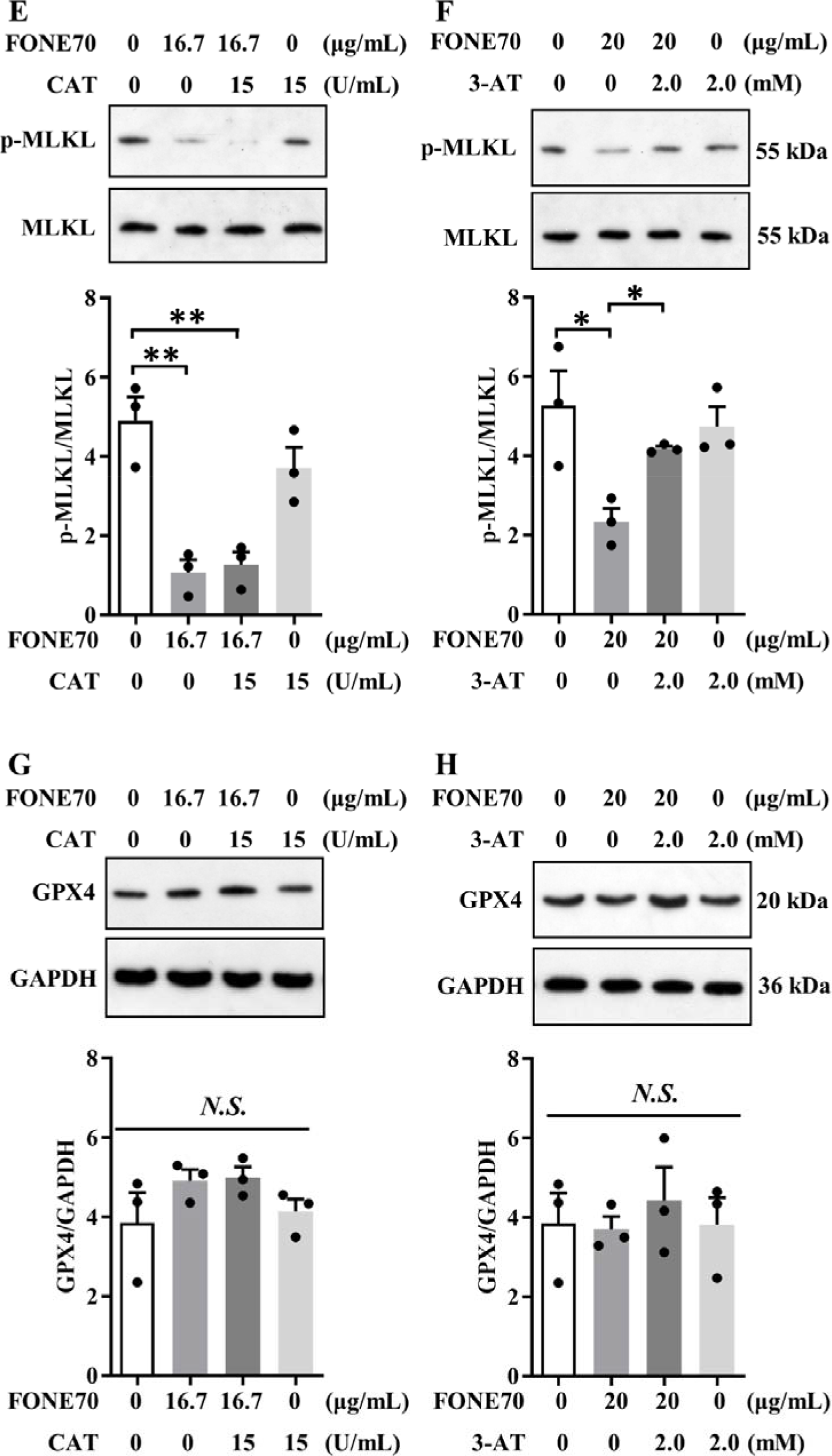
Influences of FONE/catalase-induced ROS on the protein expression of LC3B-I, LC3B-II, Cleaved Caspase 3, Pro-Caspase 3, MLKL, phospho-MLKL, GPX4, and GAPDH in macrophage Raw 264.7 cells. The cells were incubated with FONE70 at 16.7 μg/mL for 24 h in the presence of catalase (0 mM, 15 U/mL) before the expression cellular biomarkers were determined: (A) the ratio of LC3B-II/LC3B-I; (C) the ratio of Cleaved Capase 3/Pro-Caspase 3; (E) the ratio of phospho-MLKL/MLKL; (G) the ratio of GPX4/GAPDH. The cells were incubated with FONE70 at 20 μg/mL for 24 h in the presence of catalase-inhibitor 3-AT (0 mM, 2.0 mM) before the expression of cellular biomarkers were determined: (B) the ratio of LC3B-II/LC3B-I, (D) the ratio of Cleaved Capase 3/Pro-Caspase 3, (F) the ratio of phospho-MLKL/MLKL, and (H) the ratio of GPX4/GAPDH. Data were expressed as the mean ± SEM (*n* = 3). Data were analysed by one-way analysis of variance (ANOVA) followed by the Tukey’s *t*-test for post-hoc analysis, \*\**P* < 0.01, \*\**P* < 0.01, *N.S*., no significance.

It is known that ROS could induce both necrosis and apoptosis via activation of Caspase-3 protease, which is cleaved to a p12 subunit and a p17 subunit^38^. As is shown in Fig. 7C&D, the FONE70 decreased the cleaved caspase-3/pro-caspase 3 ratio (*P* < 0.01), implying the caspase-3 was not activated. The exogenous catalase didn’t resume the ratio, while the inhibition of endogenous catalase with 2 mM 3-AT successfully resumed the ratio but only to the level of normal cells. The results demonstrate the FONE-induced macrophage necrosis/ferroptosis is not caspase-3 dependent.

MLKL can cause programmed necrotic cell death, after being phosphorylated by RIP3, by binding to plasma and intracellular membranes and directly disrupting membrane integrity^39^. FONE caused plasma membrane leakage, evidenced by the rising extracellular LDH level. The p-MLKL/MLKL ratio was determined to verify whether the membrane disruption caused by FONE involves the p-MLKL. FONE70 alone decreased the p-MLKL/MLKL ratio (*P* < 0.01), indicating MLKL wasn’t phosphorylated. The addition of exogenous catalase didn’t resume the ratio, while the catalase inhibitor 3-AT managed to resume the ratio but only to the level of normal cells (Fig. 7E&F). The results elucidate that MLKL-dependent membrane disruption and necroptosis was not the cause of FONE-CAT-induced cell death.

The deactivation of glutathione peroxidase 4 (GPX4) and subsequent elevated intracellular ROS and lipid peroxidation has been recognized as one of the ferroptosis mechanisms^40^. As shown in Fig. 7G&7H, FONE70, either in the presence or absence of exogenous catalase, did not cause any notable changes in GPX4 expression. It suggests that the FONE-CAT induced ferroptosis does not involve suppressing glutathione peroxidase expression, echoing the possibility that catalase catalysed superoxide eruption directly contributed to the rising intracellular ROS.

## DISCUSSION

The addition of catalase in the polysorbate 80-stabilized fish oil nano-emulsion triggered a superoxide eruption, whose intensity was correlated to the catalase activity and the PUFAs content. Hydroperoxide in FONE was not the substrate of catalase as its content remained unchanged. Catalase is a heme-protein containing ferric iron in the resting state and has been demonstrated to conduct direct electron transfer^41^. Interaction of catalase heme with a strong reducing substrate and molecular oxygen facilitates the formation of a compound II-like intermediate containing oxyferryl heme (Fe^4+^=O), possibly catalyses the production of superoxide^17^. The enzyme, catalase, will then be reduced back to the initiate heme-(Fe^3+^) state via one-electron transfer reaction^17,42,43^. On the debate of the source of transferable electron on the lipid/water interface, S. Pullanchery et al. have recently demonstrated the ‘unexpected strong charge-transfer interactions arose from interfacial C-H⋯O hydrogen bonds’ ^44^. This provides not only an explicit explanation to the negative ζ-potential of FONE droplets, but also a possible strong electron donor exclusively available in oil emulsions. Thus, we suppose catalase promotes the electron transfer from alkyl, alkoxyl, carboxyl groups of the lipids to the neighbouring oxygen^45^ in lipid/water interfacial space and triggers the dramatic increase in superoxide (as shown in Fig. 8). Superoxide is of great importance with crucial biological and physiological roles in diverse biological systems. It is generated in the largest quantity in mitochondria as a by-product of aerobic metabolism^46^, and it is also produced by phagocytes to kill invading microorganism^47^. In addition to those elucidated sources of superoxide in biological systems, i.e., catalysation by xanthine oxidase^48^, catalase-triggered superoxide eruption in FONE constitutes a previously unrecognised mechanism. Its biological implications can be alarmingly significant, given both intracellular and extracellular ROS eruptions induced by CAT-FONE were fatal to macrophages as is explicitly shown in this study (as shown in Fig. 8).

**Fig. 8.**
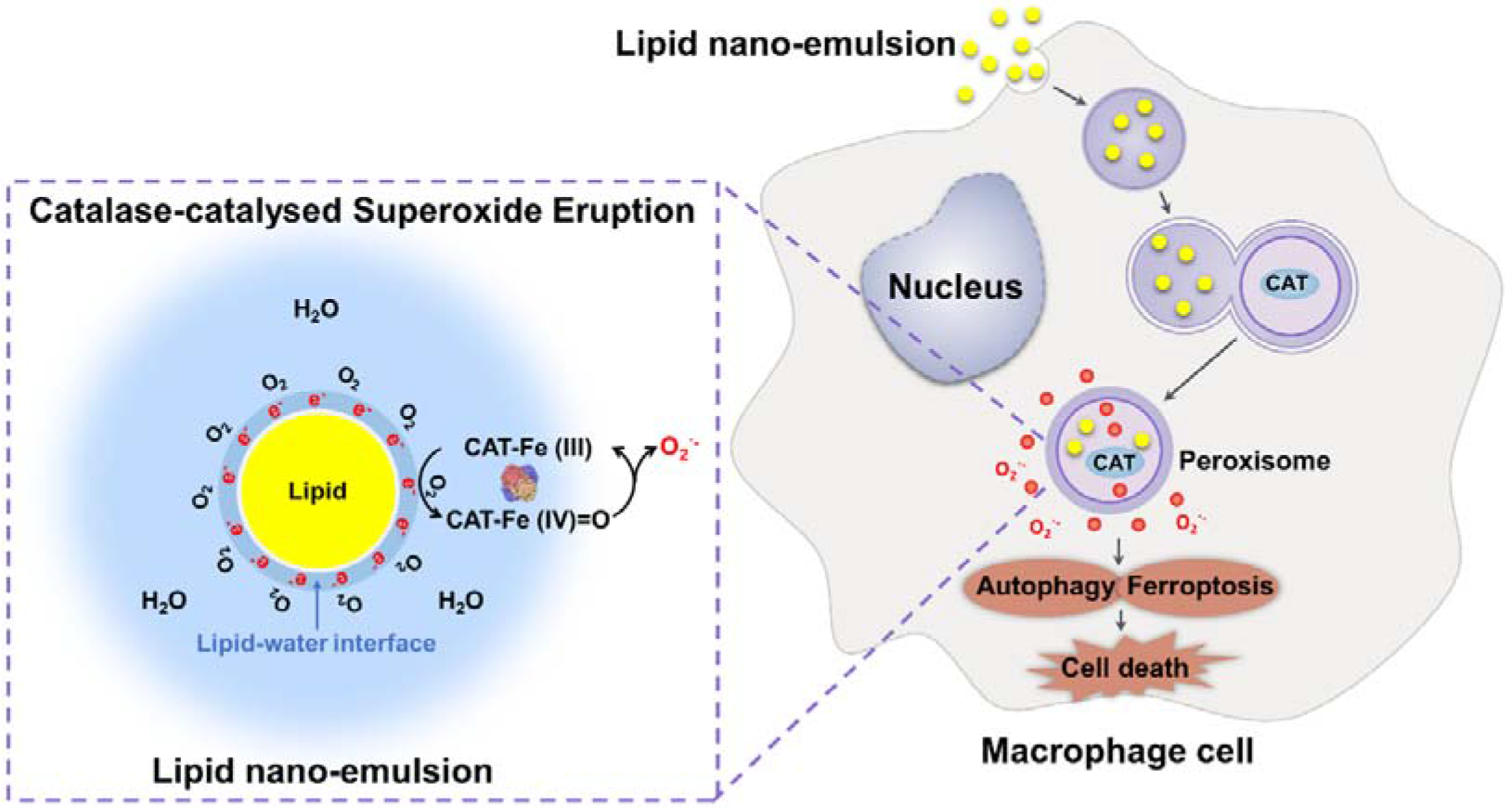
Schema of the proposed mechanism of catalase-catalysed superoxide eruption and subsequent cell death in macrophages.

To the broader extent, lipid droplets not only are produced in the engineered oil-in-water emulsions, but also exist prevalently in natural food and processed food^49^. Moreover, in a living biological system, food lipid droplets can be in touch with various tissues from the mucosal layer along the alimentary^50^ to the digestive and metabolic process involving the absorption, transportation, storage, mobilization and utilization of lipids^51^. On the other hand, catalase is available both intracellularly and extracellularly all over the mammals’ body^14,52^, which makes the encounter of lipid droplets with catalase inevitable. The resulting superoxide eruption could be a cause of inflammation on-site or even at a systemic level. As the first line of cells in the alimentary canal encountering food lipid droplets, the epithelial cells in oral mucosa and enterocytes in intestinal mucosa would be under the oxidative challenges induced by endogenous catalase, just as what happened to the macrophages. The subsequent oxidative cellular damage could play a role in the inflammatory syndromes in the alimentary tract and the irritated immune responses throughout the body.

The association of high fat-diet with both local and systemic inflammation has been extensively discussed^53,54^ but remains obscure on the mechanism of a fat-initiated cascade of the immune response. The catalase-triggered superoxide eruption reported here may provide a concise answer: the dietary fat, often emulsified by artificial emulsifiers or bile salts, would have created an enormous oil-water interface being exposed to ubiquitously distributed exogenous or endogenous catalase and resulted in superoxide eruption, which may, in turn, regulates phagocytosis and inflammation. A similar hypothesis can be drawn to explain the association of red meat consumption with gastrointestinal inflammation ^55,56^, given the high contents of fat and haemoglobin in red meat. It may also imply a new lead to understanding the side-effects of parenteral nutrition, a specialized form of food for medical use, in which PUFAs is often used in a significant amount.

The regulatory effects of PUFAs on ROS production, viability, and function of macrophages from various animals have been reported. None of the PUFAs shows cytotoxicity on the same type of murine macrophages as employed in this study, at the concentration ranging from 0.1-100 μM^57^. EPA potentiated the lipopolysaccharide (LPS)-stimulated ROS production in the macrophages, while other PUFAs, including DHA decreased ROS and reactive nitrogen species (RNS) production of the stimulated cells. At the higher fatty acid concentration, however, the viability of the head-kidney macrophages isolated from a fish was significantly decreased after 24 h or 36 h incubation with 1000 μM EPA or DHA. EPA significantly promoted the intracellular superoxide anion synthesis, which was significantly reduced by DHA^58^. In another *in vitro* study using a different murine macrophage cell line, J774, a large increase in the intracellular ROS was observed by incubation with DHA, EPA, oleic acid, linoleic acid and arachidonic acid at 100 μM for 1 h^59^, while no cytotoxicity was recorded. Furthermore, the fish oil-derived polyunsaturated *n*-3 fatty acids (e.g., EPA, DPA) were found to induce the death of mice bone marrow-derived macrophages at 200 μM through promoting intracellular ROS products as early as 30 mins after treatment^60^. In general, EPA boosts cellular ROS, despite the different origin of the macrophage cells or different treatment times. At relatively high dosages, e.g., 1000 μM (self-assembled droplets) and 200 μM (emulsion prepared by BSA-assisted sonication), EPA could induce cell death. It is consistent with the present study, where the cell death was induced with PUFA-rich fish oil emulsion, although at a much lower dosage (approx. 50–70 μM) which tempts one to speculate that emulsification may potentiate EPA’s cytotoxicity in macrophages. The higher ROS level and more severe cell damage caused by FONE70 could be well explained by the tripled content of EPA.

Another important conclusion we are proposing here is that CAT-triggered superoxide eruption in FONE can cause macrophage cell death through ferroptosis and autophagy, but not apoptosis. The cell death was effectively prohibited by using the ferroptosis inhibitor liproxstatin-1, which acts as a potent radical-trapping antioxidant^33^ to scavenge the superoxide anions. Together with the unchanged expression level of glutathione peroxidase in cells after FONE treatments, the results support the hypothesis that the superoxide rather than the unsaturated fatty acids is the direct cause of membrane lipid peroxidation and cell deformation. Moreover, the fact that cellular ROS, ferroptosis, and autophagy were all prohibited by CAT-specific inhibitor 3-AT elucidates the imperative role the CAT has played, not as an antioxidant enzyme but an oxidase to catalyse the production of superoxide and subsequent ROS. Intriguingly, EPA and DHA are known to reduce intracellular ROS and attenuate oxidative stress-induced DNA damage in vascular endothelial cells ^61^. Contrary to macrophage, vascular endothelial cells are lack CAT but high in SOD ^62^, implying that much less intracellular ROS would be produced when these cells encountered the *ω*-3 PUFA droplets. Similar reasoning may provide a new perspective to understand the nature of EPA-induced apoptosis in HepG2 human hepatoma cells ^63^, which has high CAT activity, but not in L-02 liver cells expressing less CAT.

Several possible co-factors should be put in consideration when accessing the biological impacts of the PUFA-rich emulsion in real-life setting. Firstly, the positively charged metal ions in food or body fluids may mask the negative charge of lipid droplets and block the catalase-triggered electron transfer. Secondly, the type of emulsifiers or droplet stabilisers, particularly in the case of Pickering emulsions, may alter the surface electrostatic status. It will either promote or suppress the generation of superoxide and affect the superoxide production as we have found in fish oil emulsions prepared with other emulsifiers (data not shown). Thirdly, the co-existing antioxidants and heme-containing agents other than catalase in dietary constituents or body fluids. For instance, bile salts are surfactants with potent reducing power and therefore would interfere with the oil/water interface and ROS reactions. If the emulsion was intravenously infused, it would be in company with plenty of antioxidant molecules in the blood, i.e., SOD, which instantly scavenges any superoxide produced by catalase. As the catalase’s oxidase activity has broad substrate specificity, some dietary constituents such as ferulic acid, indole-3-carbinol, and epigallocatechin-3-gallate are the enzyme’s substrates and capable of interfering the oxidation ^17^. Lastly, the influences of chain length and saturation level of fatty acids on the CAT-lipid interaction may also be just as decisive as other factors.

## CONCLUSION

Superoxide eruption from the PUFAs-rich fish oil nano-droplets was found to be catalase-initiated and -dependent in both acellular and cellular systems. The resulting superoxide anions increased cytosolic ROS and membrane lipid peroxidation, induced severe cellular oxidative damages leading to autophagy and ferroptosis. The autophagy involves activation of microtubule-associated protein LC3B. The ferroptosis was independent to protease Caspase-3 activation or glutathione peroxidase suppression. The cellular damages were significantly inhibited by the catalase-specific inhibitor, 3-Amino-1,2,4-triazole. Our findings discover a hidden risk factor of the widely acclaimed fish oil emulsion and suggest a novel mechanism of cellular damage by dietary lipids on a mucosal layer of the alimentary tract. It warrants further studies to understand the catalytic mechanism of catalase triggered superoxide production in PUFAs emulsions and to identify other cofactors affecting the cellular redox homeostasis.

## Funding

This study was supported by the National Key Research and Development Program of China (2016YFD0400202).

## Conflict of interest

The authors declare no competing financial interest.

## Notes

### Competing Interest Statement

The authors have declared no competing interest.

