## Supplemental information for "Fish oil Nano-emulsion Kills Macrophage: Ferroptosis and Autophagy Triggered by Catalase-catalysed Superoxide Eruption"

Table S1. The fatty acid content of F30 and F70 fish oil.

| **Category** | **Fatty acid** | **F30(%)*** | | **F70(%)*** | |
| --- | --- | --- | --- | --- | --- |
| SFA | Octanoic acid (C8:0) | N.D. | 24.0 | N.D. | 1.7 |
|  | Decanoic acid (C10:0) | N.D. |  | N.D. |  |
|  | Undecanoic acid (C11:0) | N.D. |  | N.D. |  |
|  | Lauric acid (C12:0) | 0.18±0.01 |  | N.D. |  |
|  | Tridecanoic acid methyl ester (C13:0) | 0.12±0.01 |  | N.D. |  |
|  | Myristic acid (C14:0) | 8.40±0.14 |  | 0.15±0.01 |  |
|  | Pentadecanoic acid (C15:0) | 0.63±0.01 |  | N.D. |  |
|  | Palmitic acid (C16:0) | 8.93±0.16 |  | 0.27±0.00 |  |
|  | Margaric acid (C17:0) | 0.57±0.00 |  | 0.07±0.00 |  |
|  | Stearic acid (C18:0) | 3.89±0.07 |  | 0.39±0.01 |  |
|  | Arachidic acid (C20:0) | 0.63±0.01 |  | 0.13±0.01 |  |
|  | Behenic acid (C22:0) | 0.28±0.02 |  | 0.15±0.00 |  |
|  | Tricosanoic acid (C23:0) | 0.33±0.02 |  | 0.51±0.04 |  |
| USFA | Myristoleic acid (C14:1) | 0.10±0.01 | 26.1 | N.D. | 4.4 |
|  | Palmitoleic acid (C16:1) | 11.10±0.19 |  | 0.20±0.00 |  |
|  | Elaidic acid (C18:1n9t) | N.D. |  | 0.63±0.01 |  |
|  | Oleic acid (C18:1n9c) (OA) | 11.28±0.18 |  | 0.27±0.00 |  |
|  | Eicosenoic acid (C20:1) | 2.07±0.05 |  | 0.16±0.01 |  |
|  | cis-11,14-Eicosadienoic acid (C20:2) | 0.52±0.01 |  | 3.15±0.09 |  |
|  | Erucic acid (C22:1n9) | 0.31±0.00 |  | N.D. |  |
|  | cis-13,16-docosadienoic acid (C22:2) | 0.14±0.00 |  | N.D. |  |
|  | nervonic acid (C24:1) | 0.54±0.01 |  | N.D. |  |
| Omega-3 | cis-5,8,11,14,17-Eicosapentaenoic acid (C20:5n3) (EPA) | 24.71±0.46 | 44.4 | 75.48±0.87 | 86.5 |
|  | cis-4,7,10,13,16,19-docosahexaenoic acid (C22:6n3) (DHA) | 17.80±0.31 |  | 10.54±0.13 |  |
|  | α-Linolenic acid (C18 : 3n3) (ALA) | 1.57±0.04 |  | 0.47±0.00 |  |
|  | cis-11,14,17-Eicosatrienoic acid (C20:3n3) | 0.30±0.01 |  | N.D. |  |
| Omega-6 | Linoleic acid (C18 : 2n6c) (LA) | 3.56±0.01 | 5.6 | 0.51±0.01 | 7.4 |
|  | γ-Linolenic acid (C18 : 3n6) (GLA) | 0.31±0.01 |  | 0.27±0.00 |  |
|  | cis-8,11,14-Eicosatrienoic acid (C20 : 3n6) | 0.23±0.01 |  | 0.31±0.00 |  |
|  | Arachidonic acid (C20 : 4n6) (AA) | 1.51±0.03 |  | 6.32±0.09 |  |

*: the percentage was given as the content of each fatty acid versus the total fatty acids content of the fish oil.

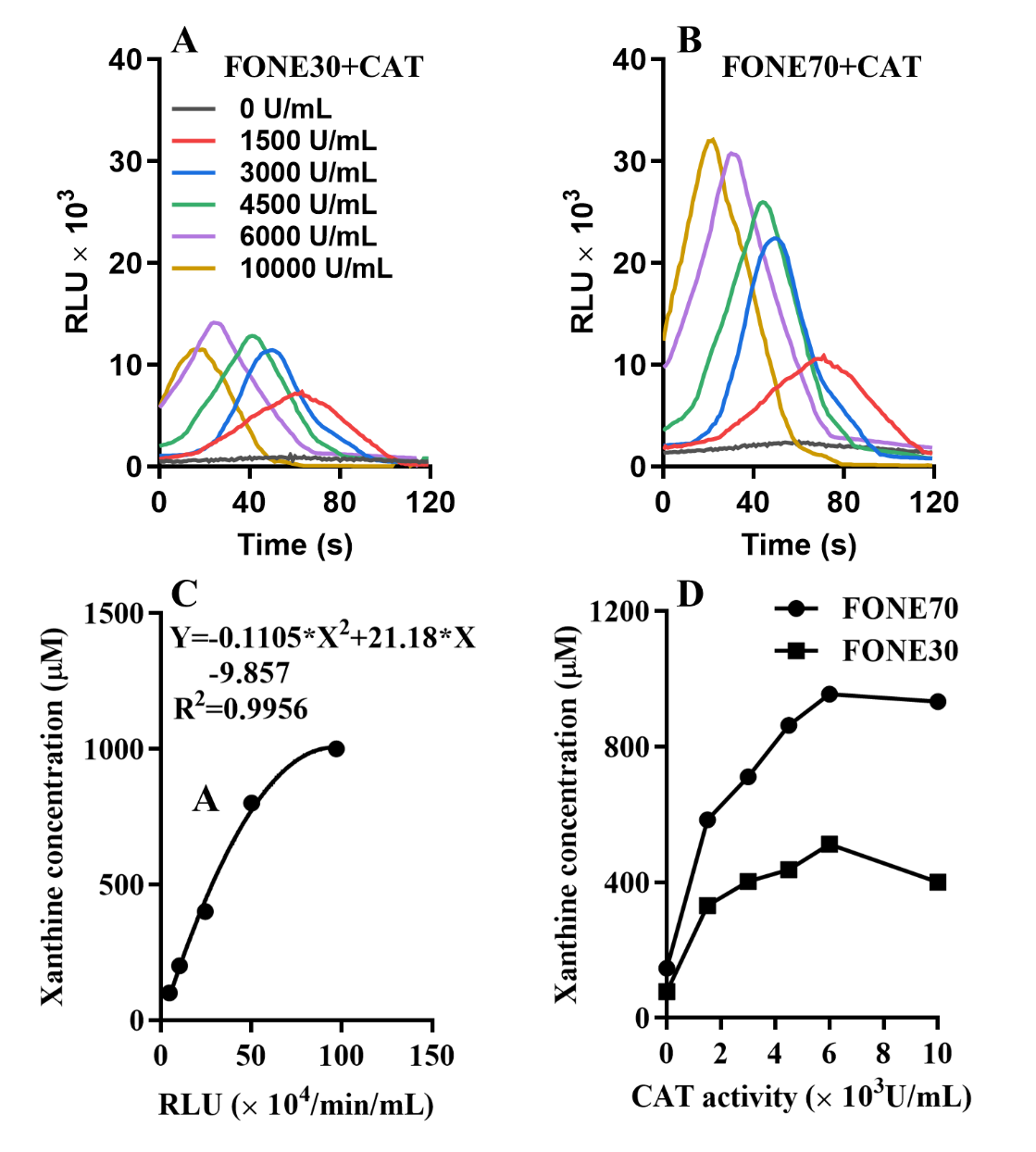

Fig. S1 Time course of chemiluminescence. (A) The effect of catalase on the time course of chemiluminescence in FONE30 using MPEC probe; (B) The effect of catalase on the time course of chemiluminescence in FONE70 using MPEC probe. (C) The standard curve of ROS was produced by Xanthine with 0.1 U/mL of Xanthine oxidase using MPEC as a probe. (D) Equivalent Xanthine concentration with 0.1 U of Xanthine oxidase of ROS in 5 mg/mL FONE30 or FONE70 in the presence of catalase at the activity of 0, 1,500, 3,000, 4,500, 6,000, 10,000 U/mL using MPEC probe. Data were expressed as the mean ± SEM (*n* = 3).

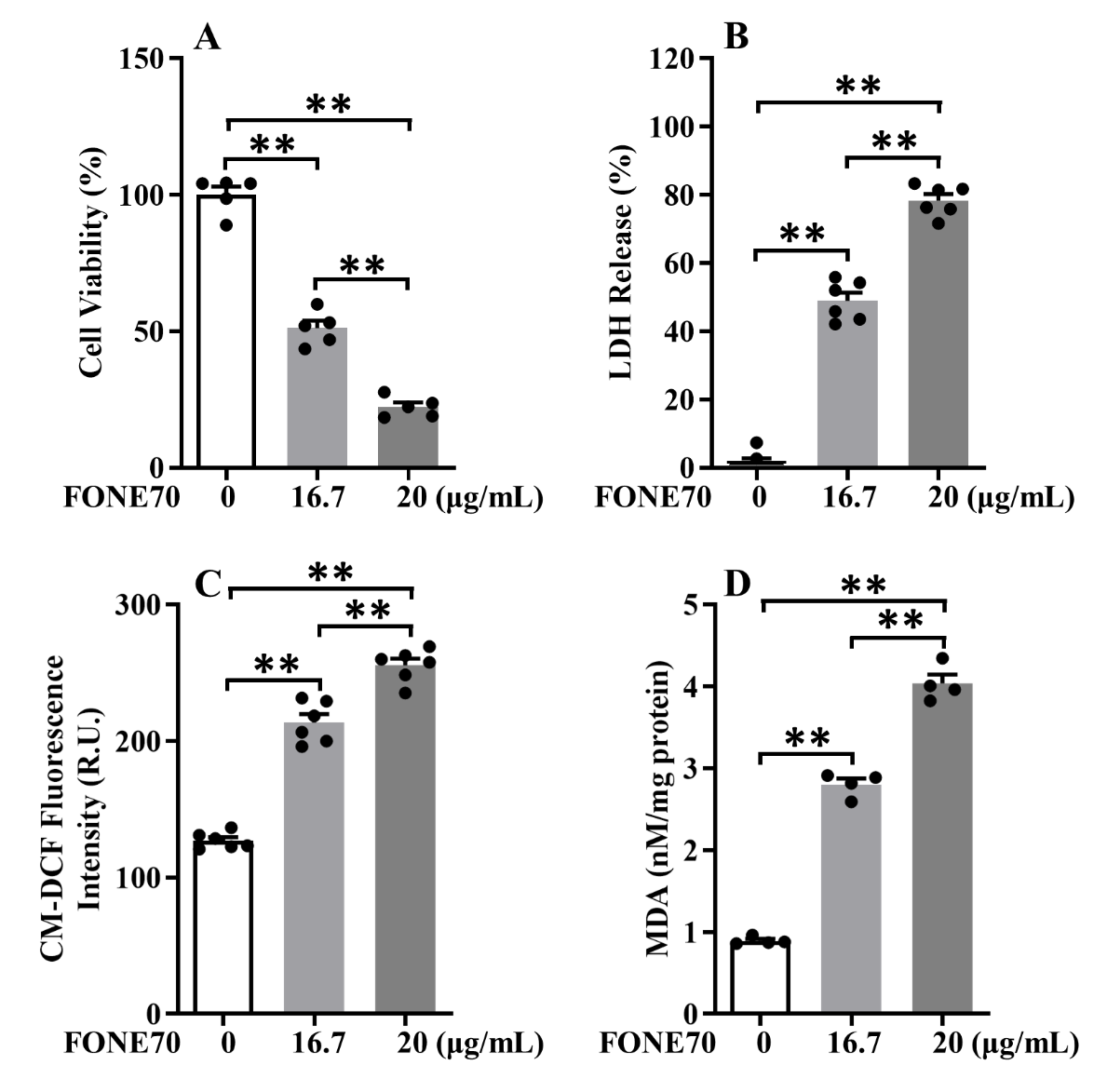

Fig. S2 The effects of FONE70 alone on Raw 264.7 cells. (A) Cell viability, (B) LDH release, (C) total ROS using a CM-H_2_DCFDA probe, (D) MDA content in cells treated with FONE70 at the concentration of 16.7 μg/mL and 20 μg/mL. Data were expressed as the mean ± SEM (*n* = 4-6). Data were analysed by one-way analysis of variance (ANOVA) followed by the Tukey’s *t*-test for post-hoc analysis, ***P* < 0.01.

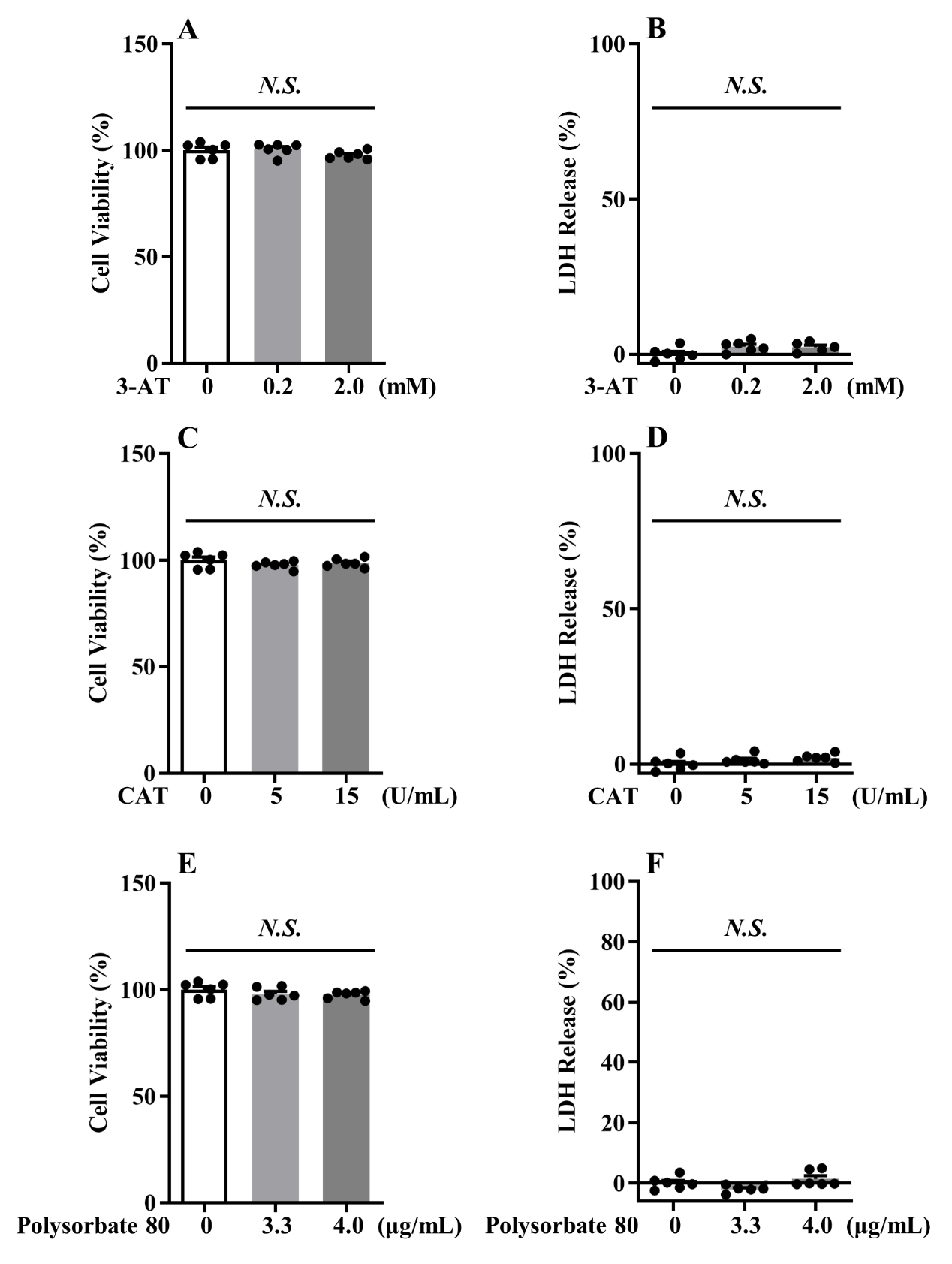

Fig. S3 Effects of 3-AT, CAT, and polysorbate 80 on the cell viability and LDH release in Raw 264.7. The effect of 3-AT on the cell viability (A) and LDH release (B) at the concentrations of 0.2 mM and 2.0 mM for 24 h in the Raw 264.7 cells. The effect of CAT on the cell viability (C) and LDH release (D) at the concentrations of 5 U/mL and 15 U/mL for 24 h in the Raw 264.7 cells. The effect of polysorbate 80 on the cell viability (E) and LDH release (F) at the concentrations of 3.3 μg/mL and 4.0 μg/mL for 24 h in the Raw 264.7 cells. Data were expressed as the mean ± SEM (*n* = 6). Data were analysed by one-way analysis of variance (ANOVA) followed by the Tukey’s *t*-test for post-hoc analysis, *N.S.*, no significance.

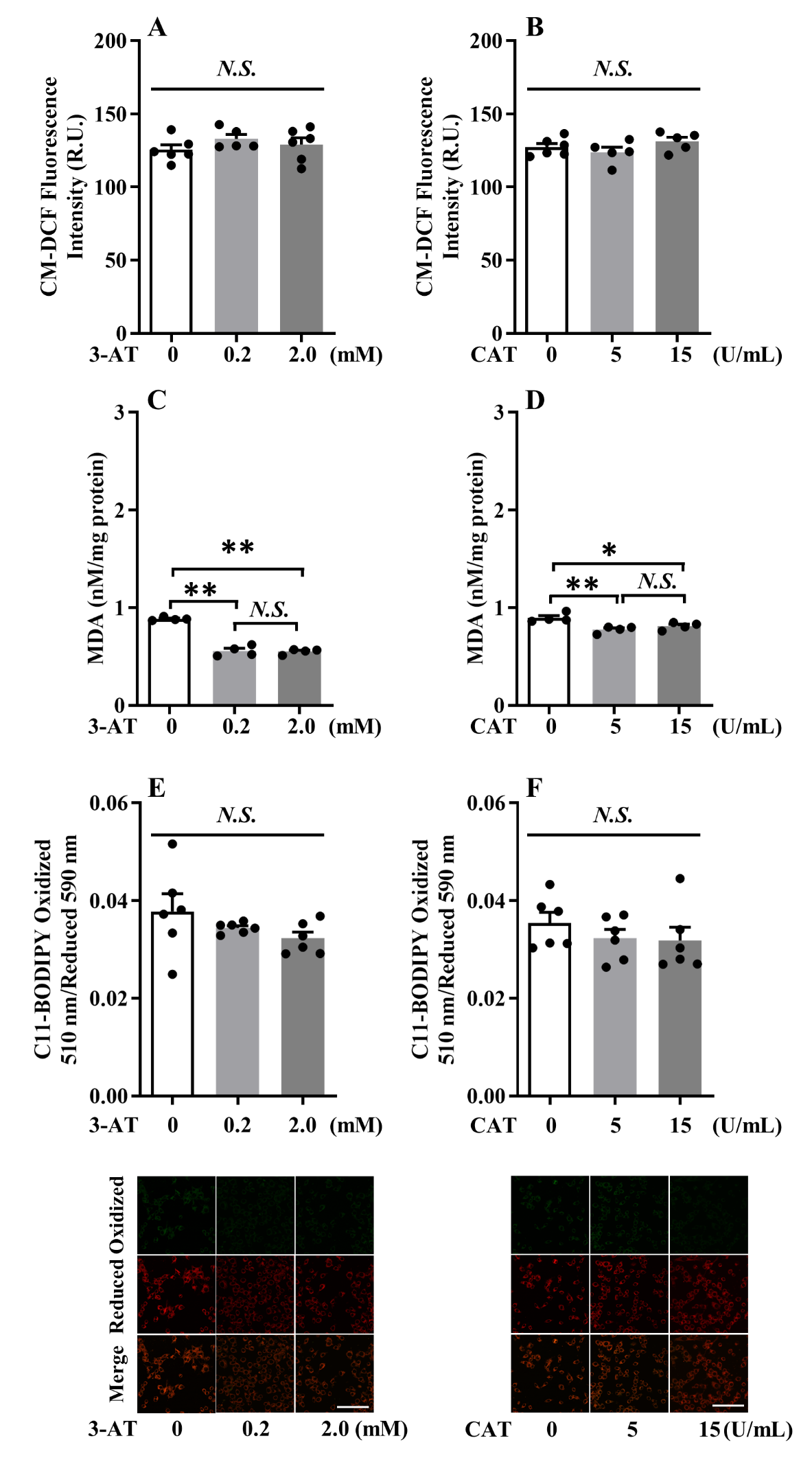

Fig. S4 Effects of 3-AT and CAT on the levels of cytosolic ROS, MDA release, and lipid peroxidation in Raw 264.7 cells. The effect of 3-AT (0.2 mM or 2.0 mM, for 24 h) on (A) the cytosolic ROS, (C) MDA Content, and (E) the ratio of oxidized 510 nm/reduced 590 nm and confocal microscopy images (scale bar = 100 μm) of lipid peroxidation using a C11-BODIPY probe in cells. The effect of CAT on (B) the levels of total ROS, (D) MDA Content, and (F) the ratio of oxidized 510 nm/reduced 590 nm and confocal microscopy images (scale bar = 100 μm) of lipid peroxidation using a C11-BODIPY probe in Raw 264.7 cells treated with CAT at the concentrations of 5 U/mL and 15 U/mL for 24 h. Data were expressed as the mean ± SEM (*n* = 4-5). Data were analysed by one-way analysis of variance (ANOVA) followed by the Tukey’s *t*-test for post-hoc analysis, **P* < 0.05, ***P* < 0.01, *N.S.*, no significance.

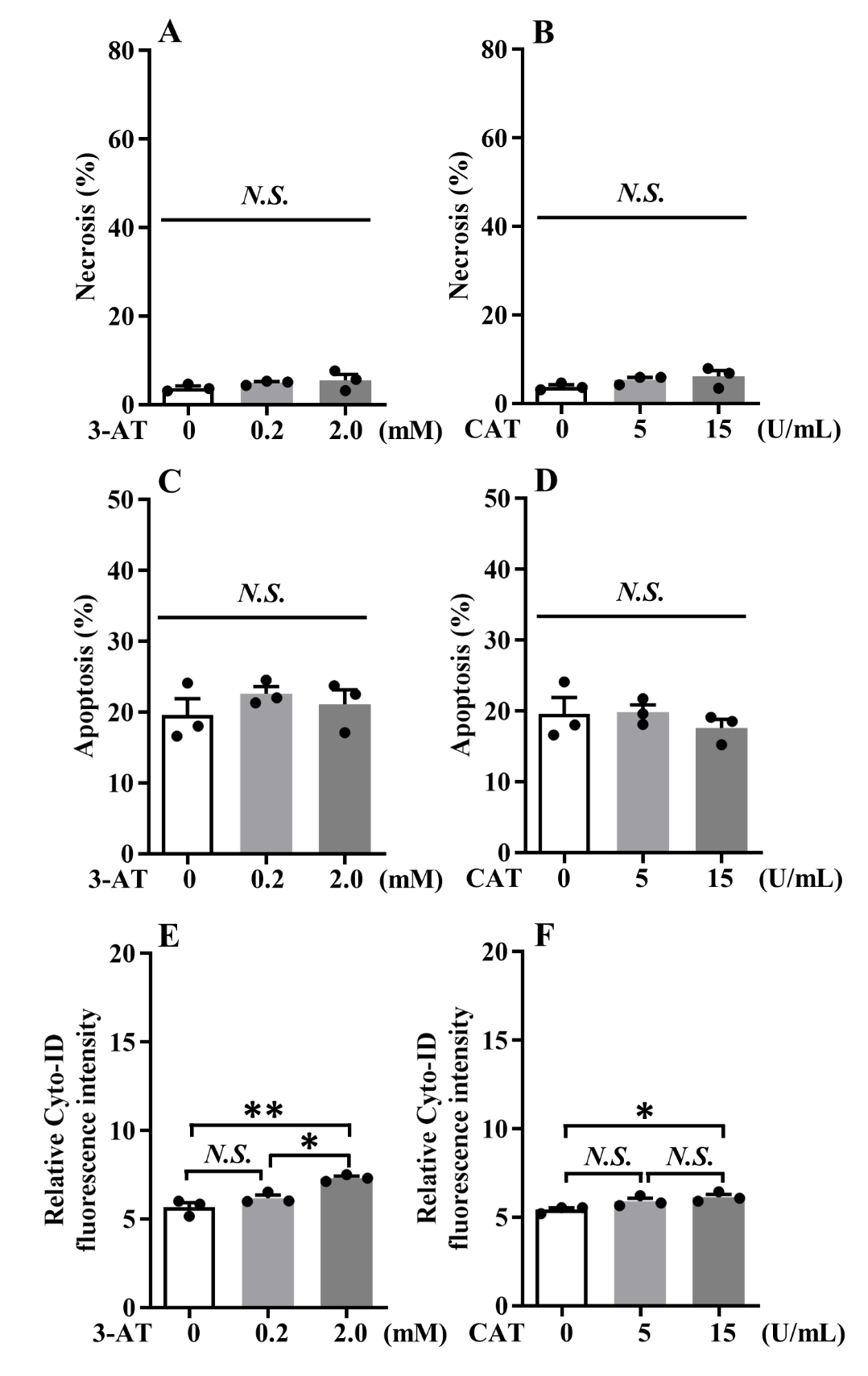

Fig. S5 Effects of 3-AT and CAT on the levels of necrosis, apoptosis, autophagic flux in Raw 264.7 cells. The effects of 3-AT (0.2 mM and 2.0 mM, for 24 h) on (A) levels of necroptosis, (C) levels of apoptosis, (E) levels of autophagy flux in the cells. The effects of CAT (5 U/mL and 15 U/mL, for 24 h) on (B) levels of necroptosis, (D) levels of apoptosis, (F) levels of autophagy flux in the cells. Data were expressed as the mean ± SEM (*n* = 3). Data were analysed by one-way analysis of variance (ANOVA) followed by the Tukey’s *t*-test for post-hoc analysis, **P* < 0.05, ***P* < 0.01, *N.S.*, no significance.
